# Mutant EZH2 alters the epigenetic network and increases epigenetic heterogeneity in B cell lymphoma

**DOI:** 10.1101/2024.08.05.606629

**Authors:** Ofir Griess, Noa Furth, Nofar Harpaz, Nicoletta Di Bernardo, Tomer-Meir Salame, Bareket Dassa, Ioannis Karagiannidis, Yusuke Isshiki, Menachem Gross, Ari M. Melnick, Wendy Béguelin, Guy Ron, Efrat Shema

## Abstract

Diffuse large B cell lymphomas and follicular lymphomas show recurrent mutations in epigenetic regulators; among these are loss-of-function mutations in KMT2D and gain-of-function mutations in EZH2. To systematically explore the effects of these mutations on the wiring of the epigenetic network, we applied a single-cell approach to probe a wide array of histone modifications. We show that mutant-EZH2 elicits extensive effects on the epigenome of lymphomas, beyond alterations to H3K27 methylations, and is dominant over KMT2D mutations. Utilizing the single-cell data, we present computational methods to measure epigenetic heterogeneity. We identify an unexpected characteristic of mutant-EZH2, but not KMT2D, in increasing heterogeneity, shedding light on a novel oncogenic mechanism mediated by this mutation. Finally, we present tools to reconstruct known interactions within the epigenetic network, as well as reveal potential novel cross talk between various modifications, validated by functional perturbations. Our work highlights novel roles for mutant-EZH2 in lymphomagenesis and establishes new concepts for measuring epigenetic heterogeneity and intra-chromatin connectivity in cancer cells.

## Introduction

Diffuse large B cell lymphoma (DLBCL) is the most common type of non-Hodgkin’s lymphoma (NHL) ^1,2^. Most of these cancers arise from germinal centers (GC), an immune compartment governed and maintained by the transcriptional repressor BCL6 ^3–5^. GCs are transient structures in which B cells are rapidly dividing and undergoing somatic hypermutation and class switch recombination of their immunoglobulin loci, rendering them inherently susceptible to malignant transformation ^6,7^. Under normal conditions, the GC reaction is tightly controlled to ensure that GC B cells stop proliferating to differentiate and form high-affinity antibody-secreting cells. Genetic lesions (mainly 3q27 translocations) at the master regulator BCL6 gene locus result in constitutive expression of this gene ^8^, ultimately leading to B cells hyperplasia and lymphomagenesis ^5,9–12^.

Epigenetic deregulation is one of the hallmarks of cancer, and specifically DLBCL ^13–15^. The chromatin landscape of cells, consisting of nucleosomes structures composed of DNA wrapped around an octamer of core histones, is decorated by an extensive array of post-transcriptional modifications (acetylation, methylation, phosphorylation etc.) ^16,17^. Precise regulation of these epigenetic modifications, which is governed by cell-type specific transcription factors (TFs) and chromatin regulators, is critical in driving differentiation programs and maintaining cell identity ^18,19^. In DLBCL, 85% of patients carry at least one mutation in prominent regulators of the chromatin landscape ^6^. The two genes that show the highest mutation frequency, as a single or double mutations, are EZH2 (20-30% of patients), which methylates histone 3 lysine 27 (H3K27me1/2/3), and KMT2D (30% of patients), which deposits mono-methylation of H3K4 (H3K4me1) ^6,14,20–24^.

EZH2 is the catalytic unit of the Polycomb Repressive Complex 2 (PRC2), associated with transcriptional repression ^25,26^. Normally, upon GC entry, B cells exhibit upregulation of EZH2, and it was shown to cooperate with BCL6 in GC formation ^15,27,28^. Germinal center B cell subtype of DLBCL (GCB-DLBCL), as well as follicular lymphomas (FL) which are the second most common type of NHL, exhibit recurrent heterozygous somatic point mutations in EZH2. These mutations converge to a single amino acid substitution, Y646, that is positioned in the catalytic methyltransferase SET domain ^29^. This gain-of-function (GOF) mutation alters EZH2 substrate preference from unmethylated lysine 27 to mono- and mainly di-methylated lysine 27 (H3K27me1/2), leading to accumulation of H3K27me3 at the expense of H3K27me1/2 ^30,31^. EZH2 GOF mutations were shown to promote lymphoid transformation and contribute to lymphoma cells’ proliferation and survival ^27,28,32^. These mutations alter epigenetic regulation, by changing the topology and function of chromatin domains, inducing spreading of H3K27me3, and silencing of Polycomb target genes ^27,28,33,34^. Moreover, mutant-EZH2 was shown to maintain the GC phenotype in DLBCL, potentially by mediating a significant increase in bivalent nucleosomes, marked by both the repressive H3K27me3 and the active tri-methylation of lysine 4 on histone H3 (H3K4me3) ^27,28,35^. These bivalent chromatin domains mark critical GC B cell promoters, perhaps maintaining them in a repressed state.

KMT2D mutations, like EZH2 GOF, were also shown to enhance B cell proliferation and promote lymphomagenesis in mice. While loss of KMT2D was suggested to affect locus-specific H3K4 methylation and dampen the expression of tumor-suppressor genes, whether it affects global levels of H3K4me1 remains controversial ^23,24^. A recent study identified interaction between KMT2D and CREBBP, providing explanation for the frequent co-selection of inactivating both chromatin regulators in lymphoma ^36^. Intriguingly, DLBCL patients also exhibit recurrent loss of KMT2D in combination with EZH2 GOF ^20,37^, yet the composite effects of these mutations on the epigenetic network and the cancer phenotype are not well understood.

DLBCL cells often carry mutations in several epigenetic regulators, rendering it challenging to isolate the effects of each mutation on the epigenome. In addition, accumulative evidence suggests extensive cross-talk and inter-relationships between various histone modifications ^16,17,38–40^. This underscores the need to evaluate mutations in epigenetic regulators as potentially affecting the entire epigenetic network rather than a single histone modification. Here, we apply a single-cell approach, based on Cytometry by Time of Flight (CyTOF), to systematically explore the effects of EZH2 GOF and KMT2D mutations, alone or in combination, on a wide panel of epigenetic modifications. Our data reveal extensive and dominant effects of mutant-EZH2 on the epigenetic and transcriptional network, and an uncharacterized oncogenic function for this mutation in increasing epigenetic heterogeneity. Interestingly, we show that the single-cell data can be leveraged to reconstruct and expose new putative interactions within the epigenetic network. Finally, we demonstrate a non-linear relationship between H3K27me3 and the lymphoma master regulator BCL6. Overall, this work highlights the power of single-cell epigenetic analysis to reveal novel principles of epigenetic deregulation in cancer.

## Results

### Single-cell epigenetic analysis reveals systematic alterations to various histone modifications mediated by EZH2 and KMT2D mutations

To systematically explore the effects of EZH2 Y646N GOF and KMT2D mutations on the epigenetic network, we employed a single-cell technology we recently developed that uses CyTOF to profile an extensive array of histone modifications in single cells ^38^ (Figure 1A). The epigenetic panel contained metal-conjugated antibodies targeting 16 histone modifications, three core histone proteins, cell cycle indicators, the proliferation marker Ki67 and the germinal center master regulator BCL6. We first analyzed two patient-derived xenografts (PDXs) and cell lines of GCB-DLBCL tumors, which either carry the Y646N mutant EZH2 allele in combination with KMT2D biallelic loss-of-function mutations (the PDX^MUT^ and the cell line OCI-Ly1) or carry two WT alleles for both genes (PDX^WT^ and the cell line OCI-Ly7). For each cell, the measurements of histone modifications were normalized to the levels of the core histones (see ‘Methods’), followed by dimensionality reduction using uniform manifold approximation and projection (UMAP) analysis. Both EZH2-mutant PDX and cell line clustered separately from the cells carrying WT EZH2, indicating differential epigenetic patterns in each line (Figure 1B-C and S1A-B). Consistent with the known functions of EZH2 mutation, cells carrying it exhibited elevated levels of H3K27me3 concomitant with a reduction in H3K27me2 (Figure 1C-D and S1B-C). While KMT2D mutations had no effect on global H3K4me1 levels in the PDXs, the OCI-Ly1 cells exhibited reduced levels of this modification in comparison to OCI-ly7 cells (Figure S1D-E). Finally, in both KMT2D-EZH2-mutant PDX and cell line we observed reduced levels of histone acetylation (Figure 1C-D and S1B-E). As these cells carry an additional mutation in the histone acetyltransferase CREBBP, this reduced acetylation is likely attributed to loss of CREBBP, rather than reflecting epigenetic alterations directly mediated by EZH2 or KMT2D mutations.

**Figure 1:**
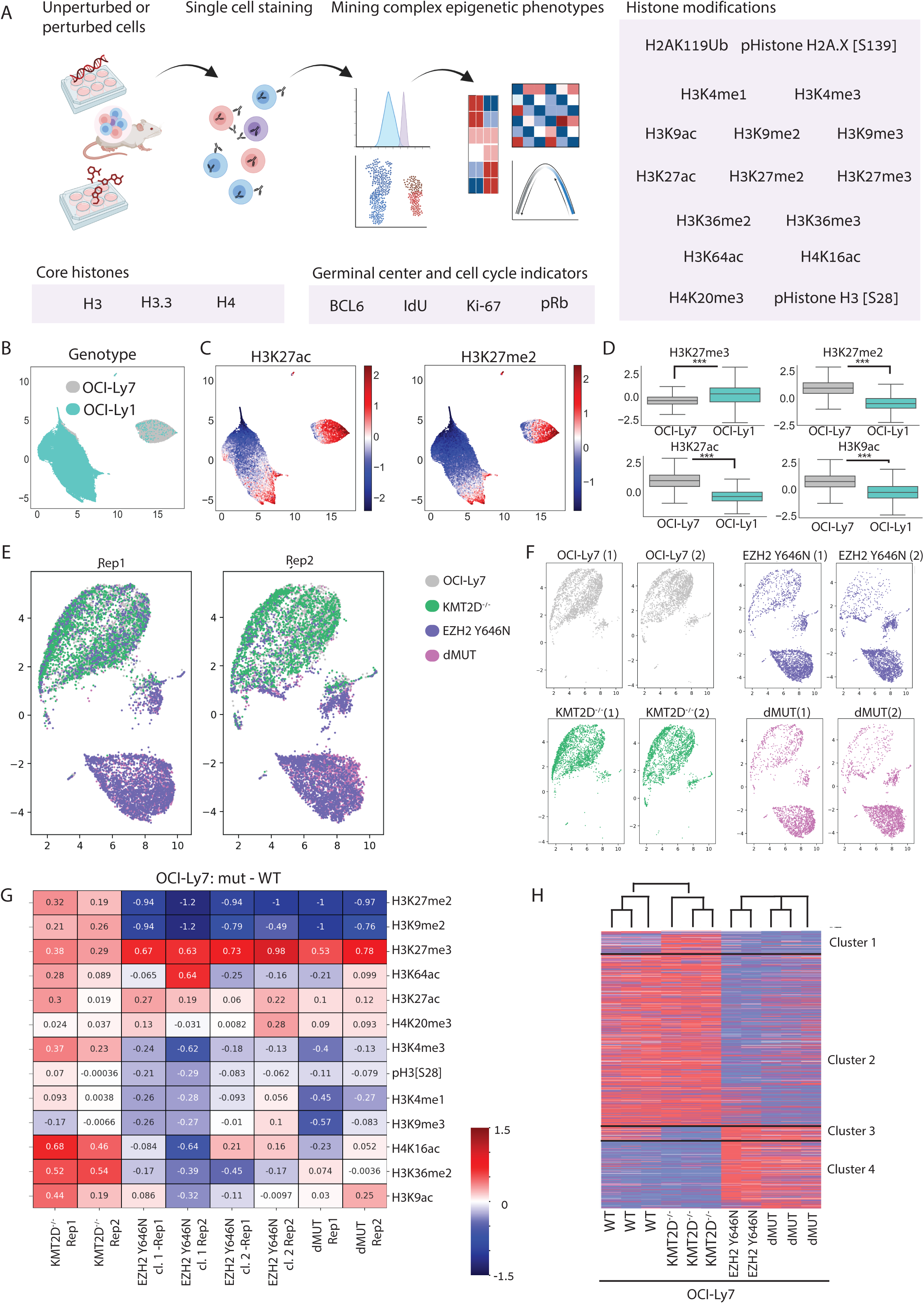
Mutant-EZH2 elicits a wide array of epigenetic alterations and shows a dominant phenotype over KMT2D mutations. **A.** Scheme of the CyTOF experimental system. Human lymphoma patient-derived xenografts, or cell lines, unperturbed or perturbed genetically or chemically, were stained with an epigenetic-oriented metal-conjugated antibody panel (see also Table S1 for list of antibodies and metals). From this single cell data, we mined complex epigenetic phenotypes exhibited by these cells. **B.** CyTOF analysis of the lymphoma patient-derived cell lines OCI-Ly7, carrying WT copies of EZH2 and KMT2D, and OCI-Ly1, carrying the EZH2 GOF mutation and biallelic loss of KMT2D. Colors indicate the sample index. **C.** Scaled, normalized levels of the indicated histone modifications on the UMAP of the cell lines that is shown in B. **D.** Expression levels of the indicated modifications in OCI-Ly1 and OCI-Ly7 cell lines, as measured by the CyTOF. P values were calculated by Welch’s t test. ***p value < 0.001. **E-F.** OCI-Ly7 cells (WT), as well as their isogenic counterparts carrying mutant-EZH2 (EZH2 Y646N), biallelic knockout of KMT2D (KMT2D^-/-^), or a combination of both mutations in EZH2 and KMT2D (dMUT), were analyzed by CyTOF. UMAP was performed based on all epigenetic marks measured in two independent biological repeats, following scaling and normalization. **E.** plotting of the indicated mutant cells on the joined UMAP in each of the repeats. **F.** Each line is shown separately. EZH2-mutant cells (alone or in combination with KMT2D loss) cluster together and separately from the WT cells, indicating a dominant epigenetic phenotype for EZH2 GOF mutation. **G.** Heatmap showing the differences in the means of the indicated histone modifications in each of the indicated mutant lines compared to the WT OCI-Ly7 cells. Two biological CyTOF replicates are shown. Red: higher in the mutant cells compared to WT. Blue: lower in mutant cells. **H.** RNA-sequencing analysis of OCI-Ly7 cells and the indicated isogenic mutant lines. Heatmap shows the rld values of differentially expressed genes between the indicated mutant lines versus OCI-Ly7. Rows are standardized and clustered; red and blue denotes high and low levels of the modifications, respectively. The list of differentially expressed genes was obtained using DESeq2 and filtered for >1 or <-1 LogFC and p.adj <= 0.05. K-means clustering = 2 (Table SX?).

To isolate the direct effects of EZH2 and KMT2D mutations, we utilized the patient-derived lymphoma cell line OCI-Ly7, which carry WT alleles of all the main epigenetic regulators that are known to be recurrently mutated in DLBCL^20,21,37^. We then generated in these OCI-Ly7 cells the EZH2 Y646N mutation, KMT2D biallelic loss-of-function mutation, or the combination of both mutations (‘dMUT’ for double-mutants), using CRISPR/Cas9 genome editing (Figure S2A-C). CyTOF analysis of the parental OCI-Ly7 cells and the different mutants revealed wide-spread epigenetic alterations mediated by EZH2 GOF mutation; while KMT2D-null cells clustered with the WT cells, most EZH2-mutant cells clustered separately (Figure 1E-F). Importantly, two independent clones of EZH2 GOF showed highly similar epigenetic phenotypes (Figure 1G). Interestingly, EZH2 GOF mutation showed a robust and dominant phenotype over the KMT2D mutation, as visualized by the highly similar epigenetic patterns of the single EZH2 Y646N mutation and the double mutant containing both EZH2 Y646N and KMT2D loss. Of note, in the double mutant the GOF mutation was generated on the background of the KMT2D null cells, thus it represents an independent transfection and a third mutant clone. This further highlights the robust and dominant effects of EZH2 GOF on mediating these epigenetic alterations. All mutants were analyzed by CyTOF in two biological repeats; cells from both repeats overlapped in the UMAP analysis and showed similar epigenetic patterns, confirming the quantitative nature of CyTOF in providing accurate measurements of histone modifications in hundreds of thousands of cells (Figure 1E and 2SD).

As expected, introducing mutant-EZH2 in this isogenic system resulted in elevated levels of H3K27me3 and reduction in H3K27me2, comparable to the levels observed in the EZH2-mutant line OCI-Ly1 (Figure 1G and S2C-D). Interestingly, all cells expressing mutant-EZH2 showed robust loss of H3K9me2 (also seen in OCI-Ly1 cells, Figure S1E), similar in extent to the loss of H3K27me2, which is the direct substrate of mutant-EZH2 enzymatic activity. These cells also showed reduced H3K9me3 levels, yet to a lesser extent, and a slight increase in H3K27ac. The isogenic system revealed that the EZH2 GOF mutation, in addition to generating a more repressed chromatin by increasing H3K27me3, also promoted loss of the ‘close’ histone H3K9 methylations, perhaps as a counter mechanism for the general repression. Interestingly, while loss of KMT2D had no effect on H3K4me1 global levels, it led to elevated levels of histone acetylation and a mild gain of H3K27me3, reminiscent of the EZH2 GOF mutants. Taken together, the data reveal a dominant role for mutant EZH2 in altering a wide array of histone modifications, beyond lysine 27 methylations, and indicate a potential cross talk between various epigenetic marks in this network.

We next aimed to explore whether these epigenetic phenotypes would translate into gene expression changes. To that end, we performed RNA sequencing of the WT OCI-Ly7 cells as well as the isogenic mutant lines (Figure 1H and S2E-H). In agreement with the mild epigenetic changes, loss of KMT2D resulted in minimal transcriptional changes (cluster 1 and 3, corresponding to genes upregulated or downregulated in KMT2D^-/-^ cells, respectively). Expression of EZH2 GOF mutation, however, led to robust transcriptional changes, with 702 genes downregulated (cluster 2) and 346 upregulated (cluster 4); the higher number of downregulated genes agrees with the function of H3K27me3 in transcriptional repression. Genomic profiling by Cut&Run in the control Ly7 cells revealed that these cluster 2 genes, presumably repressed by EZH2 GOF, were associated with higher H3K27me3 levels downstream to the TSS compared to the upregulated genes, perhaps indicating their potential regulation by this mark (Figure S2I). The independent clone of the double mutant of KMT2D^-/-^ and EZH2 Y646N showed highly similar transcriptional phenotypes to that of the single EZH2 Y646N mutation (Figure 1H and S2E). These results indicate a dominant role for mutant-EZH2 in mediating both transcriptional changes as well as epigenetic alterations.

Finally, exploring the list of up- and down-regulated genes revealed genes with interesting functions related to lymphomagenesis. For example, EZH2 mutation suppressed the expression of several genes that encode the linker histone H1 (Figure S2J). Recent studies showed that recurrent mutations in H1 genes, leading to reduced H1 levels, result in disruption of the 3D chromatin architecture and promote lymphomagenesis ^41,42^. Thus, suppression of H1 expression by mutant EZH2 may benefit lymphoma cells and lead to a more aggressive phenotype. Furthermore, both EZH2 Y646N and the double mutant cells showed upregulation of B-Cell Receptor (BCR) and CDKN1A, reported to promote survival in germinal-center B cells ^43,44^. We also observed downregulation of IL7R, which is associated with poorer prognosis in DLBCL ^45^.

### Epigenetic CyTOF reveals a role for mutant-EZH2 in inducing epigenetic variability between cells to generate higher heterogeneity

The power of the CyTOF approach is that not only does it reveal global epigenetic changes in the population due to perturbations (as seen in Figure 1), but it also enables single-cell analysis of parameters such as the variability between cells and the heterogeneity in the population. To explore these concepts, we first calculated for each epigenetic mark the Gini coefficient, commonly used to measure the inequality among the values of a frequency distribution ^46^. Interestingly, across most modifications the Gini index was higher in the EZH2-mutant cells compared to the WT, suggesting increased epigenetic heterogeneity (Figure S3A).

We next sought to define additional parameters that would reflect epigenetic heterogeneity in a population, leveraging all measurements rather than examining each modification separately. Thus, we generated a joint UMAP, based on all epigenetic modifications, of the OCI-Ly7 WT cells versus each of the isogenic mutants (Figure 2A). Heterogeneity following dimensionality reduction could potentially be manifested in different aspects (Figure 2B). We could consider global heterogeneity, as measured by (i) the global area each population of cells occupies (i.e., the area enclosed by the bounding convex hull of all points in the distribution, Figure 2Bi); and (ii) the global distance between each two cells in the population, defined by the 95th quantile of the distribution of distances between any two points in the distribution (Figure 2Bii). Another meaningful value is the local heterogeneity, reflecting whether cells in each cluster are relatively homogenous and form a tight cluster, or show greater variability to form a dispersed cluster. This can be measured by the area of the latent space occupied by the cells in a population (Figure 2Biii). Strikingly, in all these parameters, EZH2-mutant cells exhibited higher local and global epigenetic heterogeneity compared to their WT counterparts, reproducible in all experimental repeats, and irrespectively of the UMAP parameters that were used (Figure 2C and S3B). This higher heterogeneity was observed in both the EZH2 Y646N single mutant clones, as well as in the third independent clone of cells comprising the double mutant (EZH2 Y646N + KMT2D^-/-^), but was not seen in cells carrying only KMT2D biallelic loss. Finally, we also calculated the nearest-neighbor distance (NND), which refers to the distance between each point and its nearest neighbor (Figure 2Biv). If a population is more “clumped”, the distribution function for the NND will peak at shorter distances. Plotting the ratio of the cumulative distribution functions (CDFs) of EZH2-mutant cells versus WT cells revealed values below 1, indicating higher heterogeneity (corresponding to higher NND values) in EZH2-mutant cells (Figure 2D).

**Figure 2:**
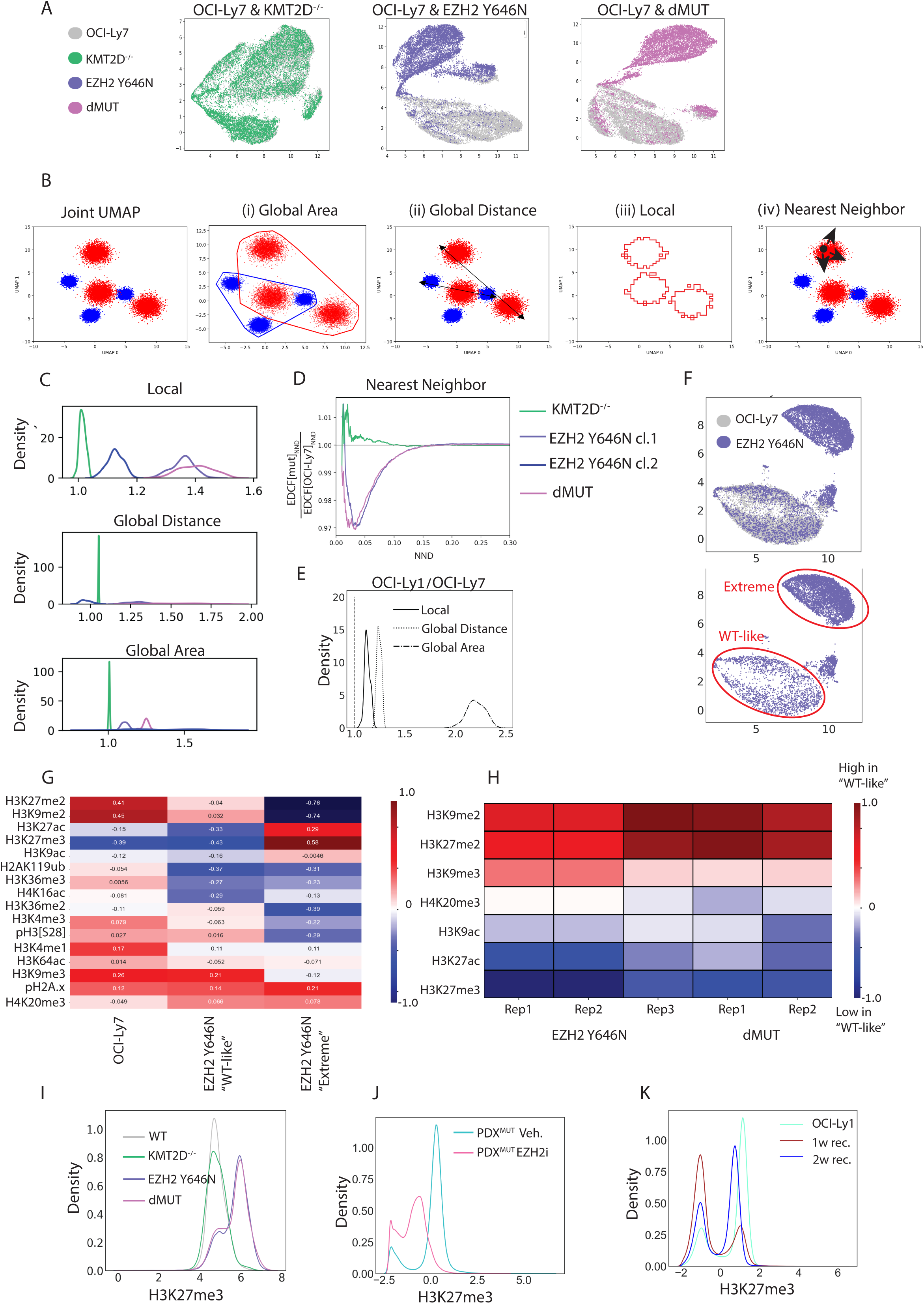
EZH2 Y646N mutation contributes to higher epigenetic heterogeneity. **A.** OCI-Ly7 cells and the indicated isogenic mutants (KMT2D^-/-^, EZH2 Y646N, and the double mutant ‘dMUT’) were analyzed by CyTOF. Joint UMAP was performed for the WT OCI-Ly7 cells versus each of the three mutants, based on all epigenetic marks measured, following scaling and normalization. Colors indicate the sample index. **B.** Illustration of the various measurements of epigenetic heterogeneity. Shown is a hypothetical UMAP of two populations, marked in red and blue. The global heterogeneity of each population is reflected by: (i) Global area, as measured by the area enclosed by the bounding convex hull of all points in the distribution, and (ii) Global distance, measured by the distance between each two cells in the population, defined by the 95th quantile of the distribution of distances between any two cells. The local heterogeneity (iii) is measured by the net area of the clusters. If clusters are more dispersed, they would occupy a larger area. (iv) Nearest-neighbor distance (NND), which refers to the distance between each point and its nearest neighbor. For a population that is more homogenous, the distribution function for the NND will peak at shorter distances. **C.** The indicated heterogeneity measurements that were defined in panel C were calculated for OCI-Ly7 cells versus each of the mutant lines, across a wide range of UMAP parameters. Plotted are the ratios of the values of each mutant line versus OCI-Ly7 (WT). Thus, values over 1 indicate excess heterogeneity in the mutant over that of the WT. EZH2 mutant cells showed higher heterogeneity in all measurements. **D.** Ratio of the cumulative distribution functions (CDFs) of the NND values in the indicated mutant lines versus the WT cells. Values below 1 indicate higher heterogeneity of EZH2-mutant cells. **E.** Heterogeneity measurements calculated as in panel C for OCI-Ly7 cells carrying WT EZH2 versus OCI-Ly1 cells expressing the mutant form. Overall, OCI-Ly1 cells carrying EZH2 GOF show higher heterogeneity. **F.** Top: Joint UMAP of OCI-Ly7 WT cells and the isogenic EZH2 Y646N cells. Colors indicate the sample index. Bottom: Only mutant-EZH2 cells are plotted, to highlight the subpopulation of mutant cells that cluster with WT cells, referred to as ‘WT-like’. The subpopulation of cells that cluster separately from WT is referred to as EZH2 Y646N ‘extreme’. **G.** Heatmap showing standardized mean values measured by CyTOF for the indicated histone modifications in OCI-Ly7 cells expressing WT EZH2 (‘WT’), and in the two populations identified in the EZH2 Y646N cells (marked in panel F), which either manifest robustly the GOF phenotype and form a distinct cluster (EZH2 Y646N ‘Extreme’), or clusters with OCI-Ly7 WT cells (EZH2 Y646N ‘WT-like’). **H.** Heatmap showing the differences in the means of the indicated histone modifications of the ‘extreme’ population compared to the ‘WT-like’ sub-population’. Shown are three biological CyTOF replicates for the single EZH2 Y646N mutant line, and two biological replicates for the double mutant. ‘WT-like’ cells show higher levels of the heterochromatin marks H3K9me2 and H3K9me3, and reduced levels of histone acetylations. **I.** Histogram of scaled and normalized H3K27me3 levels, as measured by CyTOF, in OCI-Ly7 cells (‘WT’) or the indicated isogenic mutant lines. **J.** Histogram of scaled and normalized H3K27me3 levels, as measured by CyTOF, in the PDX^MUT^ from mice treated with EZH2 inhibitor for two weeks or with vehicle as control. The two H3K27me3 subpopulations are present in both control and treated mice. EZH2 inhibition shifted the fraction of cells in each population and reduced H3K27me3 levels in the ‘high’ population. **K.** OCI-Ly1 cells expressing mutant-EZH2 were treated with EZH2 inhibitor at a concentration of 10µM for one week to completely deplete H3K27me3 (see Figure S4K). Next, the inhibitor was washed, and the cells were allowed to recover for one or two weeks. Shown is a histogram of H3K27me3 levels in untreated OCI-Ly1 cells, as well as the cells after one or two weeks of recovery. The two subpopulations of H3K27me3 are maintained throughout the experiment, yet the fraction of cells in each subpopulation shifts over time.

To further explore this phenomenon of increased heterogeneity, we examined whether it could be recapitulated in the additional EZH2-mutant systems- the PDXs and the endogenously mutant cell line OCI-Ly1. Indeed, OCI-Ly1 carrying the mutant allele EZH2 Y646N showed higher heterogeneity in all parameters compared to OCI-Ly7 (Figure 1B and 2E). Similarly, the PDX^MUT^ was more locally heterogenous than the PDX^WT^, and occupied a larger global area, although the maximal distance between cells was smaller (Figure S3C). Taken together, the analytic tools developed here indicate that EZH2 gain-of-function mutations are associated with higher epigenetic heterogeneity, perhaps contributing to the tumor’s fitness. This increased heterogeneity is unique to EZH2 mutations and is not observed in the cells containing KMT2D knockout alone, suggesting a specific characteristic of EZH2 GOF mutations rather than a general effect of mutations in any chromatin regulator.

### Mutant-EZH2 cells comprise distinct epigenetic subpopulations

Across all experiments of the isogeneic mutant-EZH2 and WT OCI-Ly7 cells, we observed that a fraction of cells expressing EZH2 Y646N clustered with the WT cells (Figure 2F and S4A-B). The percentage of these ‘WT-like’ cells was dynamic and varied between biological repeats yet was robustly detected in the two clones of the single EZH2 mutation and the double mutant with KMT2D^-/-^ (Figure S4C). These ‘WT-like’ cells seemed to reflect an intermediate epigenetic phenotype, with some modifications showing similar levels to WT cells, while others were like the EZH2-mutant cells that strongly manifest the mutation and cluster separately (EZH2 Y646N ‘Extreme’, Figure 2G and S4D). For example, while ‘WT-like’ cells had intermediate levels of H3K27me2, indicating some activity of the mutant enzyme, their H3K27me3 levels were identical to cells that did not express mutant-EZH2.

To further explore these two subpopulations within EZH2-mutant cells, we compared their epigenetic patterns across different CyTOF experiments (Figure 2H). These data revealed that ‘WT-like’ cells consistently had elevated levels of the heterochromatin marks di- and tri- methylation of histone H3 on lysine 9 (H3K9me2/3) and lower levels of H3K27ac, perhaps indicating a more compact chromatin state. While speculative, it might suggest that mutant EZH2 activity could be influenced by a pre-existing chromatin state, thus generating heterogeneity. Alternatively, these two subpopulations may reflect transitions on a differentiation axis. Importantly, these ‘WT-like’ cells were observed throughout the cell cycle, indicating they do not correspond to cells in a specific cell cycle phase (Figure S4E).

The two subpopulations comprising EZH2-mutant cells can also be seen in the bimodal distribution of H3K27me3 levels; most mutant cells exhibited elevated H3K27me3 levels, and a minor fraction of cells exhibited ‘normal’ H3K27me3 levels (Figure 2I). Of note, the H3K27me3 distribution of WT OCI-Ly7 cells or cells knocked out for KMT2D had a single peak. To examine whether this phenomenon is also present in additional patient-derived cells that endogenously carry mutant-EZH2, we analyzed the PDX^MUT^, as well as OCI-Ly1. Indeed, both models showed bimodal distribution of H3K27me3 (Figure S4F-G). These data suggest that, across different systems, EZH2-mutant cells comprise distinct subpopulations which differ in H3K27me3 levels.

We next explored whether manipulating H3K27me3 levels via inhibition of EZH2, or the H3K27me3 demethylase KDM6A/B, would affect the distribution of cells between the two subpopulations (Figure S4H). Indeed, KDM6A/B inhibition resulted in upregulation of H3K27me3, concomitant with a reduction in the fraction of ‘WT-like’ cells, from 7.1% of the population to 4.8% (Figure S4I-J). Next, we analyzed H3K27me3 single-cell distribution in the PDX^MUT^ treated in vivo for 14 days with the FDA-approved EZH2 inhibitor Tazemetostat (250 mg/kg PO, BID) ^47,48^, versus control (vehicle). Cells from Vehicle-treated mice showed bimodal distribution of H3K27me3, as expected (Figure 2J). EZH2 inhibition mainly affected the subpopulation expressing the robust GOF phenotype, reducing H3K27me3 levels in these cells. In addition, the fraction of cells comprising each of the H3K27me3 subpopulations was altered towards ‘WT-like’ cells. Importantly, both subpopulations were maintained in control and EZH2- inhibitor treated cells, suggesting this heterogeneity is an inherent feature of EZH2-mutant cancer cells. Finally, we robustly depleted H3K27me3 levels in OCI-Ly1 cells (carrying mutant-EZH2) by prolonged EZH2 inhibition, and then allowed the cells to recover for one or two weeks (Figure 2K and S4K). Interestingly, both H3K27me3 subpopulations were reestablished following a week of recovery, with time-dependent increase in the proportion of cells comprising the H3K27me3- high subpopulation (Figure 2K). These results indicate that even following a harsh perturbation, EZH2-mutant cells regenerate the two H3K27me3 subpopulations, perhaps revealing an inherent and stable feature of EZH2-mutant tumors. It highlights the power of single-cell epigenetic analysis to uncover such phenomena.

### Single-cell analysis reconstructs and reveals novel interactions within the epigenetic network

Our CyTOF data provide further support to the notion that various histone modifications are inter-connected to form a network, leading to a ‘domino effect’ of EZH2/H3K27me2/3 alterations on various additional histone marks. For each modification, cells in the population show a distribution of values, from high to low levels. We hypothesized that this information can be leveraged to gleam potential interactions between modifications, by examining coupled behaviors of epigenetic marks across cells in the population. To that end, we applied three different computational models: (1) Gradient boosting algorithm (XGBoost) that allows us to determine the effect on each modification stemming from all other modifications, calculated using the SHAP algorithm. This operation is repeated many times (>10,000) on a random subsample of the data, and the results are averaged to obtain an adjacency matrix (Figure 3A). (2) A metric we term “Probability of link”: After discretizing the data and subsampling it, we constructed a directed acyclic graph (using BNLearn - https://github.com/erdogant/bnlearn/). The operation is repeated >10,000 and the mean of each link across the adjacency matrices is calculated. Thus, this metric denotes the probability for each specific direct link to be extracted from the data (Figure 3B). (3) Partial correlations, calculated in the standard fashion; the partial correlation of modifications X and Y is the correlation of the residuals of X and Y when regressed against all other modifications (Figure 3C).

**Figure 3:**
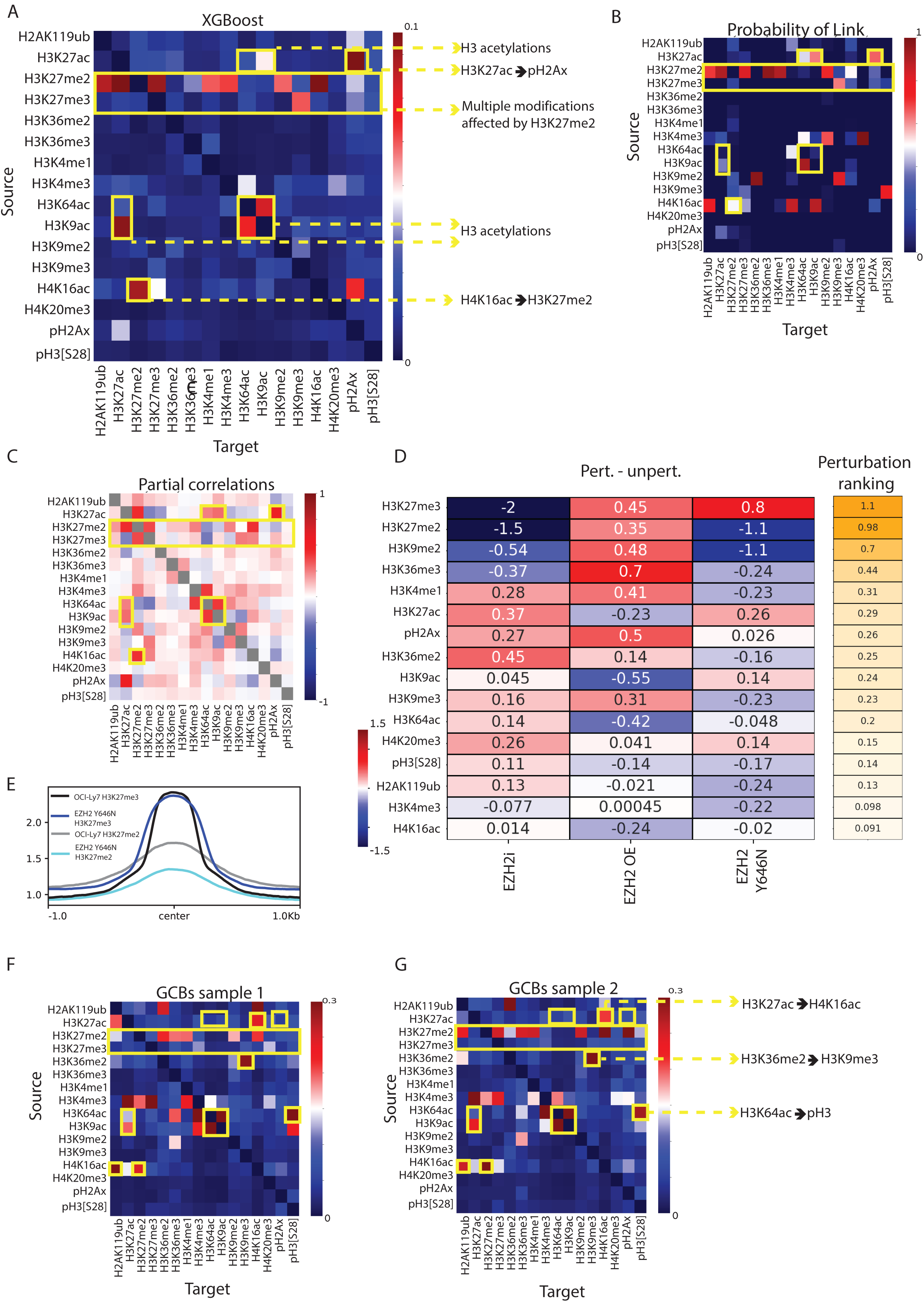
Single-cell analysis uncovers potential interactions within the epigenetic network. **A-C**. Models to decipher interactions within the epigenetic network, applied to unperturbed OCI-Ly7 single-cell CyTOF data. **A.** XGBoost analysis. **B.** ‘Probability of link’ analysis. For both A and B, the Y axis indicates ‘source’ modification and X axis indicates its ‘target’ modification affected by the Y axis. **C.** Partial correlations between histone modifications. Yellow blocks and arrows highlight connections that are discussed in the text. **D.** Left - Heatmap showing standardized mean values measured by CyTOF, for the indicated histone modifications, of unperturbed OCI-Ly7 cells compared to each of the following perturbations: EZH2 inhibition (‘EZH2i’), over expression of WT EZH2 (‘EZH2 OE’), or expression of EZH2 GOF (‘EZH2 Y646N’). For EZH2i, cells were incubated with DMSO (control) or 10µM of the EZH2 inhibitor Tazemetostat for 48h. For EZH2 OE, cells were nucleofected with PBS (control) or 1µg of plasmid expressing WT EZH2. For EZH2 Y646N, cells underwent mutagenesis for EZH2 using CRISPR/Cas9. Right – heatmap scoring the effect of the perturbation by showing the mean of the absolute change per each mark. Effect shown in descending order from high (orange) to low (white). **E.** Average coverage of Cut&Run reads of the indicated histone modifications, over H3K27me3 peaks, in OCI-Ly7 cells and the isogenic EZH2-mutant cells (EZH2 Y646N). **F-G.** XGBoost analysis done on CyTOF data of unperturbed germinal-center B cells (CD20^+^, CD38^+^), derived from tonsils, for two independent samples derived from different patients. Tonsils were dissociated to single cells followed by staining with the panel of metal-conjugated antibodies and CyTOF analysis. Yellow blocks and arrows highlight connections that are discussed in the text.

Comparing the results of the different models, we observed high overlap in reconstructing several of the well-established cross-talks between histone modifications in the network (Figure 3A-C and a biological repeat shown in S5A-C). For example, all models predicted a connection between histone H3 acetylations on lysine residues 9, 27 and 64, in agreement with their mutual deposition and removal by CREBBP/p300 and histone deacetylases, respectively. Interestingly, we found stronger connections between H3K9 and H3K64 acetylations, compared to their connections with H3K27ac; this is perhaps due to additional regulatory pathways affecting lysine 27 acetylation, such as the Polycomb complex and H3K27me2, especially in DLBCL. Another prominent example was the predicted strong effect of H3K27me2 on H3K27me3; both are deposited by EZH2, the former serves as the substrate for the latter. The network analysis also predicted an effect of H3K27ac and H4K16ac on γH2Ax; this is in-line with single-molecule studies that showed enrichment of this DNA-damage mark on nucleosomes containing H3K27ac, as well as an increase in γH2Ax in cells treated with HDAC inhibitors^35^. Moreover, a recent study reported elevated γH2Ax levels in cells carrying a deletion of MOF, the enzyme that deposits H4K16ac^49^.

An additional cross-talk reconstructed by this analysis was between H3K9 and H3K27 methylations. Previous studies reported physical and functional interactions between G9a, the enzyme that deposits H3K9me2, and EZH2^50–52^. While these studies focused on the output of these interactions on H3K27me3 deposition, our analysis predicted a connection between H3K9me2 and H3K27me2 rather than H3K27me3 (Figure 3A-C). While EZH2 is known to deposit both di- and tri-methylations, the majority of literature is focused solely on H3K27me3, due to its known regulatory function in silencing of gene expression. Surprisingly, our analyses suggest H3K27me2 as being a central node in the epigenetic network, affecting many more modifications in comparison to H3K27me3. All models predicted multiple marks affected by H3K27me2, such as H3K27ac, H2Aub, H3K36me3 and H3K4me1, while only H3K9me3 seemed to be preferentially affected by H3K27me3. H3K27ac is of particular interest, as there are multiple publications, spanning diverse biological systems, reporting a cross talk with H3K27me3^53–55^. Yet, these studies manipulated components of the Polycomb complex or lysine 27 itself, and thus cannot rule out that the effect is mediated by H3K27me2, as predicted by our data. This central role for H3K27me2 in affecting various modifications might stem from the wide distribution of this mark in the genome, as opposed to H3K27me3, which is mainly restricted to gene promoters. Finally, in addition to reconstructing known interactions, our data revealed new connections yet to be explored. For example, H4K16ac, which is also known to be widespread across large genomic regions^56^, especially in gene bodies, is predicted to affect H3K27me2 levels.

In order to experimentally validate the predicted connections, and to decouple the effects of H3K27me2 and H3K27me3, we manipulated EZH2 levels and function by three complementary means: (1) EZH2 enzymatic inhibition by Tazemetostat, leading to reduced levels of both H3K27me2 and H3K27me3; (2) Over expression of wild-type EZH2, leading to an increase in H3K27me2/3, and; (3) Expression of the gain-of-function EZH2 mutation, leading to opposing effects on these modifications: upregulation of H3K27me3 concomitant with downregulation of H3K27me2. We then scored the combined effect on all modifications in the three perturbations and ranked them in a descending order (Figure 3D). As expected, the modifications showing the largest effect were H3K27me2/3, the direct targets of EZH2. Next in ranking we observed H3K9me2, which positively correlated with the change in H3K27me2 levels rather than H3K27me3: it decreased upon EZH2 inhibition and expression of mutant-EZH2 yet increased in cells over expressing WT-EZH2. This result validates the known interaction between G9a and EZH2, in addition to providing strong support for the predicted cross-talk specifically with H3K27me2 rather than H3K27me3. H3K27ac levels also followed the change in H3K27me2, yet in an opposite direction: in both perturbations that led to loss of H3K27me2, we observed gain of H3K27ac, while in the cells gaining H3K27me2, H3K27ac levels were reduced. Finally, we also observed changes in H3K36me3 and H3K4me1, predicted to be affected by H3K27me2 in the various models. Of note, the perturbation experiments did not fully recapitulate the predicted interactions. For example, H2Aub was not affected by any of the perturbations, despite the known crosstalk between the PRC1 (which deposits H2Aub) and PRC2 complexes^57,58^, as well as the predicted link with H3K27me2 observed in our models. Taken together, these functional data support an upstream position for H3K27me2 in the epigenetic network, and validate the utility of our single-cell data and computational approaches to reveal potential coupling between modifications.

To further explore the relationship between H3 lysine 27 modifications, we profiled the genomic distributions of H3K27me3, H3K27me2 and H3K27ac in the isogenic cells expressing WT or mutant-EZH2. We first examined the full correlation matrix between all marks, to assess the extent of genomic alterations mediated by the introduction of mutant-EZH2 in the isogenic cells (Figure S5D). H3K27ac showed the highest correlation between Ly7 WT and EZH2 Y646N, as expected. Interestingly, H3K27me2 showed the highest distance between the genotypes (even compared to H3K27me3); H3K27me2 genomic distribution in EZH2-mutant cells was more like H3K27me3 compared to its patterns in the WT cells. We next analyzed the peak distribution and shape of H3K27 methylations in the isogenic cells. H3K27me3 peaks in EZH2-mutant cells were wider and showed higher basal signal across the whole genome, in agreement with spreading of this mark by the mutant enzyme (Figure 3E). Concomitantly, H3K27me2 signal was significantly reduced over these regions.

Finally, we aimed to test whether the connections observed in B cell lymphoma represent the wiring of the epigenetic network in normal germinal center B cells, or the re-wiring of this network during the tumorigenic process. To that end, we repeated the single-cell epigenetic CyTOF analysis on tonsils from two individuals, removed during tonsillectomy surgery and stained immediately without any cell culturing. The CyTOF included antibodies marking the GCB cells (CD20^+^, CD38^+^), allowing us to identify and reconstruct the network analysis on normal B cells (Figure 3F-G show XGBoost analysis of the two individuals, and S5E-F show the ‘probability of link’). Most of the connections observed in lymphoma were also seen in the normal GCB cells, including the links between histone acetylations, and the robust effect of H3K27me2 on various modifications, including H3K9me2, H3K36me3 and H3K4me1. Nevertheless, some connections were cancer-specific, such as the predicted effects of H3K27ac and H4K16ac on γH2Ax; thus, these connections in lymphoma may reflect deregulation of the DNA damage response in cancer rather than inherent wiring of the network. In addition, some new connections were observed in the normal cells and lost in lymphoma, such as a predicted effect of H3K36me2 on H3K9me3. Interestingly, all the connections were stronger in the normal cells compared to the lymphoma cells, as can be seen by comparing the heatmaps, noting the different scale (Figure 3A: scale between 0-0.1, versus Figure 3F-G: scale between 0-0.3). We speculate that this may reflect lowering of epigenetic barriers and decreased stability of the epigenetic network in cancer cells, perhaps contributing to increased heterogeneity.

### Non-linear relationship between H3K27me3 and the GC master regulator BCL6

EZH2 and BCL6 are pivotal for the formation of the germinal-center response, and deregulation of these key factors by genetic lesions in BCL6 locus and EZH2 GOF mutations promote lymphomagenesis ^9,27^. Upon perturbations of EZH2 (Figure 3D), we noticed that not only the various histone modifications were affected, but also the expression of BCL6. Counterintuitively to the expected cooperation of these two master regulators in lymphomagenesis, in our isogenic OCI-Ly7 cells, introducing EZH2 mutation led to downregulation of BCL6. This phenomenon was observed both at the protein level in our single-cell CyTOF data and by western blot (Figure 4A and S6A-B), and at the mRNA level (Figure S6C). In agreement with a potential role for H3K27me3 in suppressing BCL6 expression, Cut&Run analysis for H3K27me3 revealed gain of H3K27me3 at the BCL6 locus in cells expressing mutant-EZH2 (Figure 4B). Moreover, in the EZH2-mutant cells, H3K27me3 and BCL6 showed mild negative correlation (R=-0.16, Pearson pairwise correlation measured by CyTOF). Interestingly, in the OCI-Ly7 cells expressing WT EZH2 we observed a positive correlation between these marks (R=0.15). These seemingly contradicting observations, and the wealth of literature pointing to the importance of these factors in lymphoma, prompted us to further explore their relationship.

**Figure 4:**
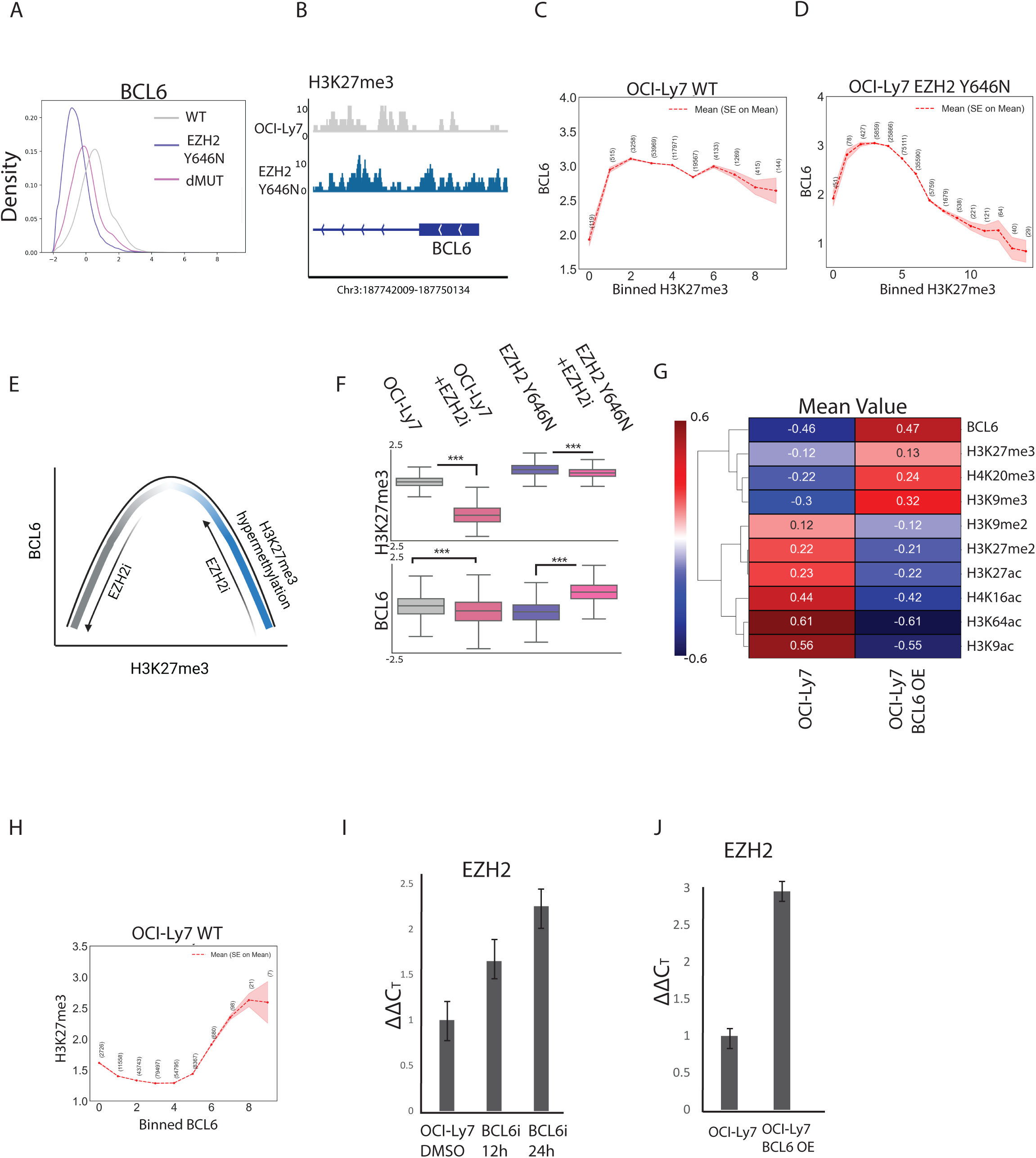
Single-cell analysis reveals non-linear relationship between H3K27me3 and BCL6. **A.** Histogram of BCL6 levels in OCI-Ly7 WT, EZH2 Y646N cells, and the double mutant of EZH2 Y646N and KMT2D^-/-^, as measured by CyTOF. **B.** Normalized coverage values of H3K27me3 on the promoter of BCL6 gene, as measured by Cut&Run, in OCI-Ly7 and EZH2 Y646N. Chr3:187742009-187750134. **C-D.** BCL6 mean levels in cells binned according to H3K27me3 levels. Red hue represents standard error of the mean. Number of cells in each bin is shown. **C.** OCI-Ly7 cells with WT EZH2. **D.** OCI-Ly7 cells with mutant-EZH2 (isogenic cell line). For this analysis measurements from four CyTOF experiments were pooled. H3K27me3 and BCL6 show a non-linear relationship. **E.** Model for the non-linear ‘bell-shaped’ relationship between H3K27me3 and BCL6. For low H3K27me3 levels, as in most OCI-Ly7 cells, it is positively correlated with BCL6. For high H3K27me3 levels, as in most EZH2-mutant cells, it is negatively correlated with BCL6. Thus, inhibiting EZH2 in the context of WT or mutant-EZH2 generates opposite effects on BCL6 levels. Generated with Biorender.com. **F.** Expression levels of H3K27me3 and BCL6, as measured by CyTOF, in the isogenic OCI-Ly7 WT and EZH2-mutant cells treated with EZH2 inhibitor for 48h at a concentration of 10µM. In OCI-Ly7 with low H3K27me3 levels, EZH2 inhibition downregulated BCL6. In the same cells expressing mutant-EZH2 and high H3K27me3, EZH2 inhibition led to upregulation of BCL6, validating a non-linear relationship. P values were calculated by Welch’s t test. ***p value < 0.001. **G.** Heatmap showing mean values of OCI-Ly7 control and BCL6 overexpressing cells, for the indicated histone modifications. BCL6 overexpression results in an increase in H3K27me3 as well as additional marks of heterochromatin, and a decrease in histone acetylation levels. **H.** H3K27me3 mean levels in cells binned according to BCL6 levels. For this analysis measurements from four CyTOF experiments were pooled. Red hue represents standard error of the mean. Number of cells in each bin is shown. **I-J.** Quantitative RT-PCR analysis of EHZ2 expression. ΔΔC_T_ values relative to OCI-Ly7 ±s.d (n=3) are shown. **I.** OCI-Ly7 cells were treated with 50µM of the BCL6 inhibitor (BCL6i) FX1 for the indicated time points. **J.** OCI-Ly7 control and BCL6 over expression. Both treatments show an increase in EZH2 expression.

We first leveraged all the data from the multiple CyTOF runs for OCI-Ly7 WT cells, binning cells according to their H3K27me3 levels and plotting the associated BCL6 levels. Interestingly, we observed a non-linear relationship: low H3K27me3 levels are positively correlated with BCL6, yet high levels are negatively correlated (Figure 4C). This non-linear ‘bell shape’ relationship was more evident in the EZH2-mutant cells, presumably because they express a wider range of H3K27me3 levels; cells expressing low H3K27me3 (such as the ‘WT-like’ cells) showed positive correlation with BCL6, while most cells expressing higher H3K27me3 showed negative correlation (Figure 4D-E). This phenomenon was recapitulated in the PDX^MUT^ and PDX^WT^ models (Figure S6D).

To further validate these non-linear dynamics, we treated the OCI-Ly7 WT and EZH2-mutant cells with EZH2 inhibitor. Confirming our single cell data, in cells expressing WT EZH2, reducing H3K27me3 by EZH2 inhibition led to downregulation of BCL6 expression, at both protein (measured by CyTOF and western blot) and RNA levels (Figure 4F and S6E-F). On the contrary, EZH2 inhibition in cells expressing mutant-EZH2 led to upregulation of BCL6, indicating a negative correlation (Figure 4F S6G-H). This upregulation was also observed in OCI- Ly1 cells carrying the EZH2 mutation (Figure S6I-J).

Finally, we tested whether manipulating BCL6 levels would affect epigenetic patterns, and specifically H3K27me3. Over expression of BCL6 in OCI-Ly7 cells led to global gain of the heterochromatin marks H3K9me3 and H4K20me3, concomitant with loss of histone acetylation, compatible with BCL6’s role as a transcriptional repressor which recruits HDAC3 to silence genes ^59^ (Figure 4G). Examining specifically the effect of BCL6 on H3K27me3 in the single-cell data, we see the expected non-liner relationship (Figure 4H). In agreement with this data, both inhibition and overexpression of BCL6 led to upregulation of EZH2 and H3K27me3 levels (Figure 4G, I-J). To conclude, our data highlights a complex non-linear relationship between BCL6 and EZH2-mediated deposition of H3K27me3. This relationship, which could only be revealed by the application of the single-cell approach, suggests a fine balance between these oncogenic pathways.

## Discussion

In this study, we present the application of CyTOF to reveal novel aspects of EZH2 gain-of-function and KMT2D loss-of-function mutations in lymphoma. EZH2 mutation was shown to alter substrate preference of EZH2 from H3K27me0 to mainly H3K27me2, manifesting in robust upregulation of H3K27me3 and loss of H3K27me2 ^31,60^. By establishing and analyzing isogenic lymphoma cells expressing WT or mutant-EZH2, we demonstrated the broad epigenetic phenotype mediated by this mutation, affecting a wide array of histone modifications other than lysine 27 methylations. For example, we observed a robust loss of H3K9me2 and gain of various histone acetylation marks in cells expressing mutant-EZH2. Moreover, the EZH2 GOF phenotype was dominant over the biallelic loss of the epigenetic regulator KMT2D; cells carrying both KMT2D knockout and mutant-EZH2 showed epigenetic phenotypes highly similar to the cells carrying only the EZH2 mutation, and were epigenetically different from KMT2D^-/-^ cells. Transcriptomic analysis revealed a similar dominant role for EZH2 Y646N mutation, indicating that the global epigenetic alterations observed by CyTOF are functionally translated into transcriptional changes. It is noteworthy that while KMT2D mutations are found across all DLBCL genetic subtypes, EZH2 mutations are unique to the GCB genetic subtypes, where they contribute to a unique disease phenotype, alluding to the dominant nature of this mutation ^21,22^.

The cross talk between the trimethylation and acetylation of lysine 27 on histone H3 (H3K27me3 and H3K27ac) is of particular interest, and was studied in diverse biological systems, from stem cells to cancer ^53,61,62^. In the OCI-Ly7 cells, introducing the genetic perturbation of EZH2 GOF mutation led to an increase in H3K27me3, which was accompanied with a mild gain of H3K27ac, suggesting a positive link between these marks. Yet, when OCI-Ly7 cells were treated with EZH2 inhibitor which reduced H3K27me3 levels, we also observed an elevation in H3K27ac levels. A similar negative correlation between H3K27me3 and H3K27ac was reported in stem cells with EZH2 knock-out, or cancer cells expressing the H3-K27M mutation, in which loss of H3K27me3 led to elevated histone acetylations ^53,61,62^. Interestingly, our network analysis, aimed at revealing direct links between histone marks, revealed that H3K27ac is tightly connected to H3K27me2, rather than H3K27me3. It is important to note that, while expression of mutant-EZH2 and application of EZH2 inhibitor exert opposite effects on H3K27me3 levels, both mediate a strong depletion of H3K27me2. Based on this data, and specifically the perturbation experiment in Figure 3D, it is tempting to speculate that the loss of H3K27me2 may be more prominent in mediating the gain of histone acetylation, rather than alterations in H3K27me3 levels. Of note, while our CyTOF panel did not include antibodies targeting the mono-methylation of histone H3 lysine 27 (H3K27me1), it was previously reported to also be downregulated in cells expressing mutant-EZH2 ^34^. This downregulation is due to the loss of mono-methylation activity by mutant-EZH2, thus generating a dependency on the WT EZH2 protein that is co-expressed in these cells, yet only from one allele. It is plausible that mono- and di-methylations of lysine 27 may function in blocking acetylation of this residue in certain genomic regions. Thus, upon H3K27me1/2 downregulation, we see accumulation of H3K27ac signal.

Epigenetic heterogeneity and plasticity have recently emerged as inherent features of cancer cells, contributing to their persistence and aggressiveness, and allowing them to cope with hostile environments such as those encountered during treatment with anti-cancer drugs ^63–65^. Yet, it remains elusive how this heterogeneity can be measured and quantified. In this work, we present several conceptual advances in leveraging single-cell epigenetic data to measure epigenetic heterogeneity. We devised computational methods to measure both local heterogeneity (i.e., within a subpopulation) and global heterogeneity (within the entire population). These methods identified a novel feature of EZH2 mutation in increasing epigenetic heterogeneity in diverse lymphoma models. Of note, loss of KMT2D did not promote heterogeneity, suggesting this is a unique feature of the EZH2 GOF mutation rather than a consequence of disruption of any epigenetic regulator. This increased heterogeneity might contribute to the oncogenic functions of mutant-EZH2, providing yet another explanation for the selective advantage it confers to the cancer cells.

One manifestation of this heterogeneity in EZH2-mutant cells is the existence of a subpopulation that, despite the GOF mutation, exhibited an attenuated phenotype and lower levels of H3K27me3. This ‘WT-like’ subpopulation was robust and maintained across different lymphoma lines, as well as *in-vivo* in the xenograft model. Given that these ‘WT-like’ cells are characterized by hypoacetylation and elevated H3K9me2/3 levels, it is possible that in these cells there is lower genomic accessibility for the mutant enzyme to act on, resulting in a milder epigenetic phenotype. It may also be that these two subpopulations of EZH2-mutant cells, expressing low/high H3K27me3, represent dynamic cell states, perhaps reflecting transitions on a differentiation axis or between light and dark zones programs. Yet another alternative possibility would be that these two subpopulations represent a dynamic balance between EZH2 and BCL6. Our single-cell data revealed a non-liner relationship between these key factors. In cells that express high BCL6, our data indicates that it represses EZH2 expression, and generates a more compact chromatin state; compatible with the phenotype of the ‘WT-like’ cells. On the contrary, cells with lower BCL6 would express higher levels of EZH2, manifesting a robust GOF phenotype. Further studies exploring the transcriptomics and probing for differentiation markers in these two subpopulations would be key in deciphering this heterogeneity, and its potential implications to lymphomagenesis.

Finally, our data strongly emphasize that perturbing a single epigenetic regulator, either by drug inhibition or by genetic manipulation, affects multiple downstream histone modifications. As various epigenetic drugs are in research use and in clinical trials for multiple cancers including lymphoma, these results highlight the need for systematic studies that view epigenetic regulation as a network, rather than focusing on a single histone mark. We demonstrate the application of several models to reveal, and potentially predict, the interactions within this network. It would be of high interest to apply these methods to single-cell epigenetic data generated in diverse biological systems, elucidating whether these interactions are conserved and can be used for tailoring better anti-cancer approaches.

## Authors’ Contributions

O.G, G.R, and E.S designed the study, performed data analysis, and wrote the manuscript. O.G and N.DB performed the experiments. N.F and N.H assisted in experimental design. T.M.S. assisted in CyTOF runs and panel design. M.G performed tonsillectomy, supplied tonsillar tissue and responsible for the IRB at Hadassah Medical Center. O.G and B.D performed bioinformatics analysis. I.K, Y.I, W.B and A. M. M designed and prepared *in-vivo* patient-derived xenografts and provided meaningful insights throughout the work on this manuscript.

## Supporting information

Figure legends for Supplementary figures

## Acknowledgements

We thank Prof. Mark Minden of the Ontario Cancer Institute, Toronto, Ontario, Canada for providing patient-derived cell lines. O.G would like to thank Dr. Anat Griess for her aid in statistical analysis and inspiring the work on DLBCL. E.S. is an incumbent of the Lisa and Jeffrey Aronin Family Career Development chair. This research was supported by grants from the European Research Council (ERC801655, ERC101115455), Emerson Collective, The Israeli Science Foundation (1881/19), Israel Cancer Research Fund, Minerva foundation, and the Israeli Council for Higher Education (CHE) via the Weizmann Data Science Research Center.

## Data availability

RNA sequencing data was submitted to GEO under the number GSE238258

Cut and Run sequencing data was submitted to GEO under the number GSE272889

Cytometry by Time of Flight data was submitted to Flow Repository under the number FR-FCM-Z6LH

## Methods

### Patient derived xenografts

Six to eight-week-old NSG mice were implanted subcutaneously in the flank with tumor fragments from a previously characterized DLBCL PDX ^66^. Tumor formation was monitored weekly by inspection and calipers. Treatment began when tumors reached 600-700 mm^3^. Mice were randomized to treatment with vehicle and Tazemetostat with 5 mice in each group. Tazemetostat was administered PO BID at 250 mg/kg in 0.5% NaCMC, 0.1% Tween-80 in dH2O by oral gavage. Mice were treated for 14 days total. Tumors were measured by caliper three times weekly. Mice were sacrificed when 14 days-treatment finished. All procedures performed were in accordance with Cornell University IACUC. For experiments in Figure 1, the untreated DLBCL PDX used were mutant EZH2^Y646N^ ^66^ and wild type EZH2 “DB-CJ” ^67^.

### Cell culture conditions and generation of transgenic lines

Cell lines (OCI-Ly1 and OCI-Ly7) were a generous gift by Dr. Mark Minden (Ontario Cancer Institute, Toronto, Ontario, Canada) and were grown in Iscove’s Modified Dulbeco Medium (IMDM) (Thermofisher: 21980032) supplemented with 20% Fetal Bovine Serum (Thermofisher: 10270106) and 1% Penicilin-Streptomycin (Biological Industries: 03-031-1B) at 37°C and 5% CO_2_. Cells were maintained at density of 0.5-2 million cells/ml. Cells were routinely checked for Mycoplasma contamination.

EZH2 Y646N mutant lines were generated through nucleofection with Lonza 4D SF kit (V4XC-2032) of 0.5 million OCI-Ly7 cells with Alt-R™ CRISPR-Cas9, sgRNA and HDR sequence (IDT). Total nucleofected population was divided into single cell colonies. DNA extraction was done with boiling of pellets in 50µL of 50mM NaOH for 10 minutes at 99°C, then supplemented with 5µL of Tris-HCl pH=8. Template was used for PCR using Hy-Tiger mix (x2) (Hylabs: EZ2031), followed by cleanup using Nucleospin Gel Extract II/PCR (Macherey-Nagel: 740609.5). 20ng clean template was analyzed by Sanger sequencing. Heterozygous clones were isolated.

BCL6 or EZH2 overexpressing cells were obtained by nucleofecting OCI-Ly7 with BCL6 overexpression plasmid pCXN2-BCL6 (Addgene: #40346) or EZH2 WT overexpression plasmid (Addgene: #173717), and compared to blank controls. Overexpression was tested using real-time PCR and Immunoblotting.

### Human subjects

All clinical studies were approved by the appropriate national research ethics committee (Helsinki committee of the Hadassah Medical Center under protocol 0058-19-HMO), and have been performed in accordance with the ethical standards.

### Tonsil processing and epigenetic profiling

Human inflamed tonsils were harvested (Helsinki committee of the Hadassah Medical Center under protocol 0058-19-HMO) and crushed into 70um mesh nylon for single cell separation. Fresh cells underwent cisplatin staining and extracellular staining as described in Cytometry by Time of Flight chapter. Cell identity profiles were done according to the tonsil atlas.^68^ To avoid batch effect of intracellular staining of histone modifications, samples were fixed in 1.6% PFA and frozen. Then all samples were thawed and stained simultaneously.

### Drug assays

1.5 million cells of OCI-Ly7 or EZH2 Y646N were incubated with DMSO, EPZ-6438 10µM (Enco: 16174-5), GSK-J4 1µM (Merck: 420205-10MG), FX1 50µM (Databiotech: S8591-5mg) for 12, 24 or 48 hours. Inhibitor effects were validated by immunoblotting for H3K27me3 (EPZ-6438 or GSK-J4) or using real-time PCR for CXCR4 (FX1).

### Immunoblotting

Whole cell lysates were prepared by suspension with 4x Laemmli sample buffer (Bio-rad: 1610747) and 50µM DTT and repeated boiling at 95°C and vortexing. Diluted 1x sample was loaded onto 4-20% Novex Tris-Glycine gel (Thermofisher: XP04205BOX) at 115V for 1h. Transfer was executed using semi dry Bio-rad Trans turbo blotter with 0.2µm nitrocellulose membranes (Bio-rad: 1704158). Then, membrane was blocked with 5% non-fat milk and washed with TBS-T. Incubation of primary antibody was conducted overnight in 5% BSA solution supplied with 0.04% sodium azide. Incubation with secondary antibody was conducted for 1h in 5% non-fat milk. Membranes were read with Clarity Western ECL Substrate (Bio-rad: 1705061). Antibodies were diluted according to the manufacturers’ instructions.

### RNA extraction and real-time PCR

RNA was extracted from 2 million cells using Nucleospin RNA isolation kit (Macherey-Nagel: 740955.50). Quality of RNA was assessed by running in agarose gel and measuring concentration in nanodrop. 500ng of total RNA was used for first strand synthesis using M-MLV Reverse transcriptase (Promega: M1701). Real-time PCR was done by using KAPA SYBR FAST ABI Prism 2X qPCR Master Mix (KAPA Biosystems: KK4603). We normalized with the housekeeping gene HPRT and calculated ΔΔCT values to evaluate differential expression.

### MARS-seq and RNA expression analysis

Total 20ng of RNA were used for library preparation according to a published protocol^69^. Three replicates per each genotype were used in the preparation. Libraries were then sequenced using a NextSeq 500/550 High Output Kit v2.5 on NextSeq500. Analysis was conducted using the previously published User-friendly Transcriptome Analysis Pipeline (UTAP)^70^. Differential expression was obtained by using DESeq2^71^. Genes showing 2-fold change expression and p.adj values of <=0.05 were considered differentially expressed.

### Cut and Run sample preparation and data analysis

Cut and Run assay was executed as described in previous works^72^. OCI-Ly7 cells were harvested, with 200,000 cells taken per reaction. Permeabilized cells, bound to Concanavalin A-coated beads, were mixed with individual primary antibody (H3K27me3: CSTC36B11, H3K27me2: abcam24684, H3K27ac: CSTD5E4) and incubated overnight at 4°C. Secondary antibody, anti-mouse HRP (JIR 115-035-003), was used as a negative control. pAG-MNase enzyme (generated in the Department of Life Sciences Core Facilities, WIS, using Addgene plasmid 123461) was added to each sample followed by incubation step of 1 h at 4°C. Targeted digestion was done by 15 min incubation on ice block (0°C) under low salt conditions. DNA purification was done using Nucleospin gel and PCR clean-up kit (Machery-Nagel, 740609). Sequencing was done on a Next-Seq 500 instrument (Illumina) using a V2 150 cycles mid output kit, allocating 10M reads per sample (paired end sequencing).

Paired-end reads of each sample (∼15.7M average reads per sample) were preprocessed with cutadapt^73^, to remove adapters and low-quality bases (parameters: --times 2 -q 30 -m 20), following evaluation of quality with FastQC. Reads were mapped to human genome (hg38, UCSC) using Bowtie^74^ version 2.4.5 (--local --very-sensitive-local --no-unal --no-mixed --no-discordant --dovetail -I 10 -X 700). Nucleosome fragments at the length >120bp were selected from the remaining unique reads using picard-tools. Alignments were normalized based on spike-in scaling factor calculated by *E. coli* “carry-over” DNA from the pAG-Mnase, as reported by Meers & Henikoff^72,75^. Bigwig files were constructed from BAM alignments using deepTools2 suite^76^, by ‘bamCoverage’ command, in 10bp bins. Heatmaps and reads coverage around TSS was visualized using missingDataAsZero parameter. Peaks were called using SEACR^75^ algorithm, with Empirical false discovery rate of 0.01.

### Antibody conjugation

Antibodies (BSA and azide free) were conjugated to metals using the MIBItag Conjugation Kit (IONpath) according to the manufacturer’s instructions.

### Cytometry by Time of Flight (CyTOF)

6 million cells were harvested and washed with Maxpar CyTOF PBS (Fluidigm: 201058). Live/dead staining was conducted by resuspension of the cells with 1.25µM Cisplatin (Fluidigm: 201064) in CyTOF PBS for 1 minute. Cisplatin was neutralized with warm IMDM supplied with 10% FBS (Invitrogen: 10270106). Fixation and permeabilization was conducted by incubation of cells with Maxpar nuclear antigen staining buffer set (Fluidigm: 201063) according to the manufacturer’s instructions. Samples were multiplexed by taking 3 million cells/sample and using the Cell-ID 20-Plex Pd Barcoding Kit (Fluidigm: 201060). After barcoding, 1 million cells from each sample were pooled to prevent batch effect and blocked using FBS for 10 minutes. Then, the pooled sample was directly incubated for 30 minutes at room temperature with the mixture of antibodies and washed with Maxpar Cell Staining Buffer (Fluidigm: 201068). Cells were fixed with fresh 4% PFA overnight. On acquisition day, samples were suspended with 4% PFA supplied with Cell-ID Intercalator-Iridium (Fluidigm: 201192A) at 125nM. Cells were then washed with Maxpar Cell acquisition solution (Fluidigm: 201244) and suspended again in Maxpar Cell acquisition solution supplied with 10% EQ Four Element Calibration Beads (Fluidigm: 201078). Prior to the run on the CyTOF machine, cells were filtered through a 35µM mesh cell strainer. Data was acquired on a Fluidigm Helios CyTOF system. The list of antibodies and conjugated metals is provided in Supplementary Table S1.

## CyTOF Data analysis

### Data processing

CyTOF data underwent the following pre-processing prior to analyses: First, the CyTOF software by Fluidigm was used for normalization and concatenation of the acquired data. Then, several gates were applied using the Cytobank platform (Beckman Coulter):

First, the normalization beads were gated out using the 140Ce channel. Then, live single cells were gated using the cisplatin 195Pt, iridium DNA label in 193Ir, event length, and the Gaussian parameters of width, center, offset and residual channels. CyTOF software was then used for samples de-barcoding ^77^.

### Data manipulation and scaling

The data analysis pipeline follows the same general scheme presented in our previous work ^38^, with some modifications. Here we outline the steps taken.

Before beginning the analysis procedure, the cells were gated using the core histones (H3, H3.3, and H4). The gate allowed only cells with a minimum raw value of 5 for all core histones into the next steps of the analysis. For the PDX lines gating was also applied on the CD45 human vs mouse markers to remove all mouse cells.

For the various markers, a hyperbolic arcsine transform (with a scale factor of 5) was first applied to the data:

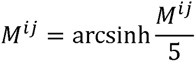

The data were then normalized to remove systematic effects using the measured values of the core histones:

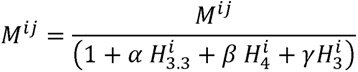

Where as in [34] the coefficients were selected to minimize the sum of the variances of the core histones, and from normalization y = (1 -*α* - *β*). A reduction of a few dozen percent in the sum of variances was observed after the subtraction, indicating the removal of systematic effects.

Finally, the data were z-transformed:

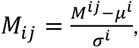

where in all the previous i denotes the observation, j denotes the column, *µ^i^* and *σ^i^* are the column mean and standard deviation, respectively. Z-transform scaling was done at the same time on samples that were acquired together.

## References

1. Alizadeh, A.A., Eisen, M.B., Davis, R.E., Ma, C., Lossos, I.S., Rosenwald, A., Boldrick, J.C., Sabet, H., Tran, T., Yu, X., et al. (2000). Distinct types of diffuse large B-cell lymphoma identified by gene expression profiling. Nature 403, 503–511. 10.1038/35000501.

2. Swerdlow, S.H., Campo, E., Pileri, S.A., Harris, N.L., Stein, H., Siebert, R., Advani, R., Ghielmini, M., Salles, G.A., Zelenetz, A.D., and Jaffe, E.S. (2016). The 2016 revision of the World Health Organization classification of lymphoid neoplasms. Blood 127, 2375–2390. 10.1182/blood-2016-01-643569.

3. Crotty, S., Johnston, R.J., and Schoenberger, S.P. (2010). Effectors and memories: Bcl-6 and Blimp-1 in T and B lymphocyte differentiation. Nat Immunol 11, 114–120. 10.1038/ni.1837.

4. Song, S., and Matthias, P.D. (2018). The Transcriptional Regulation of Germinal Center Formation. Front Immunol 9, 2026. 10.3389/fimmu.2018.02026.

5. Gatto, D., and Brink, R. (2010). The germinal center reaction. J Allergy Clin Immunol 126, 898–907; quiz 908-899. 10.1016/j.jaci.2010.09.007.

6. Pasqualucci, L., and Dalla-Favera, R. (2018). Genetics of diffuse large B-cell lymphoma. Blood 131, 2307–2319. 10.1182/blood-2017-11-764332.

7. Pasqualucci, L., Neumeister, P., Goossens, T., Nanjangud, G., Chaganti, R.S., Küppers, R., and Dalla-Favera, R. (2001). Hypermutation of multiple proto-oncogenes in B-cell diffuse large-cell lymphomas. Nature 412, 341–346. 10.1038/35085588.

8. Ye, B.H., Lista, F., Lo Coco, F., Knowles, D.M., Offit, K., Chaganti, R.S., and Dalla-Favera, R. (1993). Alterations of a zinc finger-encoding gene, BCL-6, in diffuse large-cell lymphoma. Science 262, 747–750. 10.1126/science.8235596.

9. Yang, H., and Green, M.R. (2019). Epigenetic Programing of B-Cell Lymphoma by BCL6 and Its Genetic Deregulation. Front Cell Dev Biol 7, 272. 10.3389/fcell.2019.00272.

10. Stavnezer, J., Guikema, J.E., and Schrader, C.E. (2008). Mechanism and regulation of class switch recombination. Annu Rev Immunol 26, 261–292. 10.1146/annurev.immunol.26.021607.090248.

11. Klein, U., and Dalla-Favera, R. (2008). Germinal centres: role in B-cell physiology and malignancy. Nat Rev Immunol 8, 22–33. 10.1038/nri2217.

12. Bunting, K.L., and Melnick, A.M. (2013). New effector functions and regulatory mechanisms of BCL6 in normal and malignant lymphocytes. Curr Opin Immunol 25, 339–346. 10.1016/j.coi.2013.05.003.

13. Hanahan, D. (2022). Hallmarks of Cancer: New Dimensions. Cancer Discov 12, 31–46. 10.1158/2159-8290.Cd-21-1059.

14. Jiang, Y., and Melnick, A. (2015). The epigenetic basis of diffuse large B-cell lymphoma. Semin Hematol 52, 86–96. 10.1053/j.seminhematol.2015.01.003.

15. Caganova, M., Carrisi, C., Varano, G., Mainoldi, F., Zanardi, F., Germain, P.L., George, L., Alberghini, F., Ferrarini, L., Talukder, A.K., et al. (2013). Germinal center dysregulation by histone methyltransferase EZH2 promotes lymphomagenesis. J Clin Invest 123, 5009–5022. 10.1172/jci70626.

16. Millán-Zambrano, G., Burton, A., Bannister, A.J., and Schneider, R. (2022). Histone post-translational modifications - cause and consequence of genome function. Nat Rev Genet 23, 563–580. 10.1038/s41576-022-00468-7.

17. Furth, N., and Shema, E. (2022). It’s all in the combination: decoding the epigenome for cancer research and diagnostics. Curr Opin Genet Dev 73, 101899. 10.1016/j.gde.2022.101899.

18. Shema, E., Bernstein, B.E., and Buenrostro, J.D. (2019). Single-cell and single-molecule epigenomics to uncover genome regulation at unprecedented resolution. Nat Genet 51, 19–25. 10.1038/s41588-018-0290-x.

19. Atlasi, Y., and Stunnenberg, H.G. (2017). The interplay of epigenetic marks during stem cell differentiation and development. Nat Rev Genet 18, 643–658. 10.1038/nrg.2017.57.

20. Schmitz, R., Wright, G.W., Huang, D.W., Johnson, C.A., Phelan, J.D., Wang, J.Q., Roulland, S., Kasbekar, M., Young, R.M., Shaffer, A.L., et al. (2018). Genetics and Pathogenesis of Diffuse Large B-Cell Lymphoma. N Engl J Med 378, 1396–1407. 10.1056/NEJMoa1801445.

21. Wright, G.W., Huang, D.W., Phelan, J.D., Coulibaly, Z.A., Roulland, S., Young, R.M., Wang, J.Q., Schmitz, R., Morin, R.D., Tang, J., et al. (2020). A Probabilistic Classification Tool for Genetic Subtypes of Diffuse Large B Cell Lymphoma with Therapeutic Implications. Cancer Cell 37, 551–568.e514. 10.1016/j.ccell.2020.03.015.

22. Pasqualucci, L., Trifonov, V., Fabbri, G., Ma, J., Rossi, D., Chiarenza, A., Wells, V.A., Grunn, A., Messina, M., Elliot, O., et al. (2011). Analysis of the coding genome of diffuse large B-cell lymphoma. Nat Genet 43, 830–837. 10.1038/ng.892.

23. Zhang, J., Dominguez-Sola, D., Hussein, S., Lee, J.E., Holmes, A.B., Bansal, M., Vlasevska, S., Mo, T., Tang, H., Basso, K., et al. (2015). Disruption of KMT2D perturbs germinal center B cell development and promotes lymphomagenesis. Nat Med 21, 1190–1198. 10.1038/nm.3940.

24. Ortega-Molina, A., Boss, I.W., Canela, A., Pan, H., Jiang, Y., Zhao, C., Jiang, M., Hu, D., Agirre, X., Niesvizky, I., et al. (2015). The histone lysine methyltransferase KMT2D sustains a gene expression program that represses B cell lymphoma development. Nat Med 21, 1199–1208. 10.1038/nm.3943.

25. Heyn, H., and Esteller, M. (2013). EZH2: an epigenetic gatekeeper promoting lymphomagenesis. Cancer Cell 23, 563–565. 10.1016/j.ccr.2013.04.028.

26. Comet, I., Riising, E.M., Leblanc, B., and Helin, K. (2016). Maintaining cell identity: PRC2- mediated regulation of transcription and cancer. Nat Rev Cancer 16, 803–810. 10.1038/nrc.2016.83.

27. Béguelin, W., Popovic, R., Teater, M., Jiang, Y., Bunting, K.L., Rosen, M., Shen, H., Yang, S.N., Wang, L., Ezponda, T., et al. (2013). EZH2 is required for germinal center formation and somatic EZH2 mutations promote lymphoid transformation. Cancer Cell 23, 677–692. 10.1016/j.ccr.2013.04.011.

28. Béguelin, W., Teater, M., Gearhart, M.D., Calvo Fernández, M.T., Goldstein, R.L., Cárdenas, M.G., Hatzi, K., Rosen, M., Shen, H., Corcoran, C.M., et al. (2016). EZH2 and BCL6 Cooperate to Assemble CBX8-BCOR Complex to Repress Bivalent Promoters, Mediate Germinal Center Formation and Lymphomagenesis. Cancer Cell 30, 197–213. 10.1016/j.ccell.2016.07.006.

29. Morin, R.D., Johnson, N.A., Severson, T.M., Mungall, A.J., An, J., Goya, R., Paul, J.E., Boyle, M., Woolcock, B.W., Kuchenbauer, F., et al. (2010). Somatic mutations altering EZH2 (Tyr641) in follicular and diffuse large B-cell lymphomas of germinal-center origin. Nat Genet 42, 181–185. 10.1038/ng.518.

30. McCabe, M.T., Graves, A.P., Ganji, G., Diaz, E., Halsey, W.S., Jiang, Y., Smitheman, K.N., Ott, H.M., Pappalardi, M.B., Allen, K.E., et al. (2012). Mutation of A677 in histone methyltransferase EZH2 in human B-cell lymphoma promotes hypertrimethylation of histone H3 on lysine 27 (H3K27). Proc Natl Acad Sci U S A 109, 2989–2994. 10.1073/pnas.1116418109.

31. McCabe, M.T., Ott, H.M., Ganji, G., Korenchuk, S., Thompson, C., Van Aller, G.S., Liu, Y., Graves, A.P., Della Pietra, A., 3rd, Diaz, E., et al. (2012). EZH2 inhibition as a therapeutic strategy for lymphoma with EZH2-activating mutations. Nature 492, 108–112. 10.1038/nature11606.

32. Velichutina, I., Shaknovich, R., Geng, H., Johnson, N.A., Gascoyne, R.D., Melnick, A.M., and Elemento, O. (2010). EZH2-mediated epigenetic silencing in germinal center B cells contributes to proliferation and lymphomagenesis. Blood 116, 5247–5255. 10.1182/blood-2010-04-280149.

33. Donaldson-Collier, M.C., Sungalee, S., Zufferey, M., Tavernari, D., Katanayeva, N., Battistello, E., Mina, M., Douglass, K.M., Rey, T., Raynaud, F., et al. (2019). EZH2 oncogenic mutations drive epigenetic, transcriptional, and structural changes within chromatin domains. Nat Genet 51, 517–528. 10.1038/s41588-018-0338-y.

34. Béguelin, W., Teater, M., Meydan, C., Hoehn, K.B., Phillip, J.M., Soshnev, A.A., Venturutti, L., Rivas, M.A., Calvo-Fernández, M.T., Gutierrez, J., et al. (2020). Mutant EZH2 Induces a Pre-malignant Lymphoma Niche by Reprogramming the Immune Response. Cancer Cell 37, 655–673.e611. 10.1016/j.ccell.2020.04.004.

35. Shema, E., Jones, D., Shoresh, N., Donohue, L., Ram, O., and Bernstein, B.E. (2016). Single-molecule decoding of combinatorially modified nucleosomes. Science 352, 717–721. 10.1126/science.aad7701.

36. Vlasevska, S., Garcia-Ibanez, L., Duval, R., Holmes, A.B., Jahan, R., Cai, B., Kim, A., Mo, T., Basso, K., Soni, R.K., et al. (2023). KMT2D acetylation by CREBBP reveals a cooperative functional interaction at enhancers in normal and malignant germinal center B cells. Proc Natl Acad Sci U S A 120, e2218330120. 10.1073/pnas.2218330120.

37. Chapuy, B., Stewart, C., Dunford, A.J., Kim, J., Kamburov, A., Redd, R.A., Lawrence, M.S., Roemer, M.G.M., Li, A.J., Ziepert, M., et al. (2018). Molecular subtypes of diffuse large B cell lymphoma are associated with distinct pathogenic mechanisms and outcomes. Nat Med 24, 679–690. 10.1038/s41591-018-0016-8.

38. Harpaz, N., Mittelman, T., Beresh, O., Griess, O., Furth, N., Salame, T.M., Oren, R., Fellus-Alyagor, L., Harmelin, A., Alexandrescu, S., et al. (2022). Single-cell epigenetic analysis reveals principles of chromatin states in H3.3-K27M gliomas. Mol Cell 82, 2696–2713.e2699. 10.1016/j.molcel.2022.05.023.

39. Lasserre, J., Chung, H.R., and Vingron, M. (2013). Finding associations among histone modifications using sparse partial correlation networks. PLoS Comput Biol 9, e1003168. 10.1371/journal.pcbi.1003168.

40. Perner, J., Lasserre, J., Kinkley, S., Vingron, M., and Chung, H.R. (2014). Inference of interactions between chromatin modifiers and histone modifications: from ChIP-Seq data to chromatin-signaling. Nucleic Acids Res 42, 13689–13695. 10.1093/nar/gku1234.

41. Yusufova, N., Kloetgen, A., Teater, M., Osunsade, A., Camarillo, J.M., Chin, C.R., Doane, A.S., Venters, B.J., Portillo-Ledesma, S., Conway, J., et al. (2021). Histone H1 loss drives lymphoma by disrupting 3D chromatin architecture. Nature 589, 299–305. 10.1038/s41586-020-3017-y.

42. Willcockson, M.A., Healton, S.E., Weiss, C.N., Bartholdy, B.A., Botbol, Y., Mishra, L.N., Sidhwani, D.S., Wilson, T.J., Pinto, H.B., Maron, M.I., et al. (2021). H1 histones control the epigenetic landscape by local chromatin compaction. Nature 589, 293–298. 10.1038/s41586-020-3032-z.

43. Béguelin, W., Rivas, M.A., Calvo Fernández, M.T., Teater, M., Purwada, A., Redmond, D., Shen, H., Challman, M.F., Elemento, O., Singh, A., and Melnick, A.M. (2017). EZH2 enables germinal centre formation through epigenetic silencing of CDKN1A and an Rb-E2F1 feedback loop. Nat Commun 8, 877. 10.1038/s41467-017-01029-x.

44. Young, R.M., Shaffer, A.L., 3rd, Phelan, J.D., and Staudt, L.M. (2015). B-cell receptor signaling in diffuse large B-cell lymphoma. Semin Hematol 52, 77–85. 10.1053/j.seminhematol.2015.01.008.

45. Zhao, Y., Cui, W.L., Feng, Z.Y., Xue, J., Gulinaer, A., and Zhang, W. (2020). Expression of Foxp3 and interleukin-7 receptor and clinicopathological characteristics of patients with diffuse large B-cell lymphoma. Oncol Lett 19, 2755–2764. 10.3892/ol.2020.11374.

46. Gini, C. (1936). On the measure of concentration with espacial reference to income and wealth. Cowles Commission 2.

47. Study of Tazemetostat in Participants With Relapsed or Refractory B-cell Non-Hodgkin’s Lymphoma With EZH2 Gene Mutation. https://clinicaltrials.gov/ct2/show/NCT03456726.

48. Morschhauser, F., Tilly, H., Chaidos, A., McKay, P., Phillips, T., Assouline, S., Batlevi, C.L., Campbell, P., Ribrag, V., and Damaj, G.L. (2020). Tazemetostat for patients with relapsed or refractory follicular lymphoma: an open-label, single-arm, multicentre, phase 2 trial. The Lancet Oncology 21, 1433–1442.

49. Valerio, D.G., Xu, H., Chen, C.W., Hoshii, T., Eisold, M.E., Delaney, C., Cusan, M., Deshpande, A.J., Huang, C.H., Lujambio, A., et al. (2017). Histone Acetyltransferase Activity of MOF Is Required for MLL-AF9 Leukemogenesis. Cancer Res 77, 1753–1762. 10.1158/0008-5472.Can-16-2374.

50. Mozzetta, C., Pontis, J., Fritsch, L., Robin, P., Portoso, M., Proux, C., Margueron, R., and Ait-Si-Ali, S. (2014). The histone H3 lysine 9 methyltransferases G9a and GLP regulate polycomb repressive complex 2-mediated gene silencing. Mol Cell 53, 277–289. 10.1016/j.molcel.2013.12.005.

51. Mozzetta, C., Pontis, J., and Ait-Si-Ali, S. (2015). Functional Crosstalk Between Lysine Methyltransferases on Histone Substrates: The Case of G9A/GLP and Polycomb Repressive Complex 2. Antioxid Redox Signal 22, 1365–1381. 10.1089/ars.2014.6116.

52. Casciello, F., Kelly, G.M., Ramarao-Milne, P., Kamal, N., Stewart, T.A., Mukhopadhyay, P., Kazakoff, S.H., Miranda, M., Kim, D., Davis, F.M., et al. (2022). Combined Inhibition of G9a and EZH2 Suppresses Tumor Growth via Synergistic Induction of IL24-Mediated Apoptosis. Cancer Res 82, 1208–1221. 10.1158/0008-5472.Can-21-2218.

53. Lavarone, E., Barbieri, C.M., and Pasini, D. (2019). Dissecting the role of H3K27 acetylation and methylation in PRC2 mediated control of cellular identity. Nat Commun 10, 1679. 10.1038/s41467-019-09624-w.

54. Pasini, D., Malatesta, M., Jung, H.R., Walfridsson, J., Willer, A., Olsson, L., Skotte, J., Wutz, A., Porse, B., Jensen, O.N., and Helin, K. (2010). Characterization of an antagonistic switch between histone H3 lysine 27 methylation and acetylation in the transcriptional regulation of Polycomb group target genes. Nucleic Acids Res 38, 4958–4969. 10.1093/nar/gkq244.

55. Tie, F., Banerjee, R., Stratton, C.A., Prasad-Sinha, J., Stepanik, V., Zlobin, A., Diaz, M.O., Scacheri, P.C., and Harte, P.J. (2009). CBP-mediated acetylation of histone H3 lysine 27 antagonizes Drosophila Polycomb silencing. Development 136, 3131–3141. 10.1242/dev.037127.

56. Radzisheuskaya, A., Shliaha, P.V., Grinev, V.V., Shlyueva, D., Damhofer, H., Koche, R., Gorshkov, V., Kovalchuk, S., Zhan, Y., Rodriguez, K.L., et al. (2021). Complex-dependent histone acetyltransferase activity of KAT8 determines its role in transcription and cellular homeostasis. Mol Cell 81, 1749–1765.e1748. 10.1016/j.molcel.2021.02.012.

57. Laugesen, A., Højfeldt, J.W., and Helin, K. (2019). Molecular Mechanisms Directing PRC2 Recruitment and H3K27 Methylation. Mol Cell 74, 8–18. 10.1016/j.molcel.2019.03.011.

58. Kim, J.J., and Kingston, R.E. (2022). Context-specific Polycomb mechanisms in development. Nat Rev Genet 23, 680–695. 10.1038/s41576-022-00499-0.

59. Hatzi, K., Jiang, Y., Huang, C., Garrett-Bakelman, F., Gearhart, M.D., Giannopoulou, E.G., Zumbo, P., Kirouac, K., Bhaskara, S., Polo, J.M., et al. (2013). A hybrid mechanism of action for BCL6 in B cells defined by formation of functionally distinct complexes at enhancers and promoters. Cell Rep 4, 578–588. 10.1016/j.celrep.2013.06.016.

60. Sneeringer, C.J., Scott, M.P., Kuntz, K.W., Knutson, S.K., Pollock, R.M., Richon, V.M., and Copeland, R.A. (2010). Coordinated activities of wild-type plus mutant EZH2 drive tumor-associated hypertrimethylation of lysine 27 on histone H3 (H3K27) in human B-cell lymphomas. Proc Natl Acad Sci U S A 107, 20980–20985. 10.1073/pnas.1012525107.

61. Mohammad, F., Weissmann, S., Leblanc, B., Pandey, D.P., Højfeldt, J.W., Comet, I., Zheng, C., Johansen, J.V., Rapin, N., Porse, B.T., et al. (2017). EZH2 is a potential therapeutic target for H3K27M-mutant pediatric gliomas. Nat Med 23, 483–492. 10.1038/nm.4293.

62. Piunti, A., Hashizume, R., Morgan, M.A., Bartom, E.T., Horbinski, C.M., Marshall, S.A., Rendleman, E.J., Ma, Q., Takahashi, Y.H., Woodfin, A.R., et al. (2017). Therapeutic targeting of polycomb and BET bromodomain proteins in diffuse intrinsic pontine gliomas. Nat Med 23, 493–500. 10.1038/nm.4296.

63. Flavahan, W.A., Gaskell, E., and Bernstein, B.E. (2017). Epigenetic plasticity and the hallmarks of cancer. Science 357. 10.1126/science.aal2380.

64. Feinberg, A.P., and Levchenko, A. (2023). Epigenetics as a mediator of plasticity in cancer. Science 379, eaaw3835. 10.1126/science.aaw3835.

65. De, S., Shaknovich, R., Riester, M., Elemento, O., Geng, H., Kormaksson, M., Jiang, Y., Woolcock, B., Johnson, N., Polo, J.M., et al. (2013). Aberration in DNA methylation in B-cell lymphomas has a complex origin and increases with disease severity. PLoS Genet 9, e1003137. 10.1371/journal.pgen.1003137.

66. Chapuy, B., Cheng, H., Watahiki, A., Ducar, M.D., Tan, Y., Chen, L., Roemer, M.G., Ouyang, J., Christie, A.L., Zhang, L., et al. (2016). Diffuse large B-cell lymphoma patient-derived xenograft models capture the molecular and biological heterogeneity of the disease. Blood 127, 2203–2213. 10.1182/blood-2015-09-672352.

67. Kotlov, N., Bagaev, A., Revuelta, M.V., Phillip, J.M., Cacciapuoti, M.T., Antysheva, Z., Svekolkin, V., Tikhonova, E., Miheecheva, N., Kuzkina, N., et al. (2021). Clinical and Biological Subtypes of B-cell Lymphoma Revealed by Microenvironmental Signatures. Cancer Discov 11, 1468–1489. 10.1158/2159-8290.Cd-20-0839.

68. Massoni-Badosa, R., Aguilar-Fernández, S., Nieto, J.C., Soler-Vila, P., Elosua-Bayes, M., Marchese, D., Kulis, M., Vilas-Zornoza, A., Bühler, M.M., Rashmi, S., et al. (2024). An atlas of cells in the human tonsil. Immunity 57, 379–399.e318. 10.1016/j.immuni.2024.01.006.

69. Jaitin, D.A., Kenigsberg, E., Keren-Shaul, H., Elefant, N., Paul, F., Zaretsky, I., Mildner, A., Cohen, N., Jung, S., Tanay, A., and Amit, I. (2014). Massively parallel single-cell RNA-seq for marker-free decomposition of tissues into cell types. Science 343, 776–779. 10.1126/science.1247651.

70. Kohen, R., Barlev, J., Hornung, G., Stelzer, G., Feldmesser, E., Kogan, K., Safran, M., and Leshkowitz, D. (2019). UTAP: User-friendly Transcriptome Analysis Pipeline. BMC Bioinformatics 20, 154. 10.1186/s12859-019-2728-2.

71. Love, M.I., Huber, W., and Anders, S. (2014). Moderated estimation of fold change and dispersion for RNA-seq data with DESeq2. Genome Biol 15, 550. 10.1186/s13059-014-0550-8.

72. Meers, M.P., Bryson, T.D., Henikoff, J.G., and Henikoff, S. (2019). Improved CUT&RUN chromatin profiling tools. Elife 8. 10.7554/eLife.46314.

73. Martin, M. (2011). Cutadapt removes adapter sequences from high-throughput sequencing reads. EMBnet. journal 17, 10–12.

74. Langmead, B., and Salzberg, S.L. (2012). Fast gapped-read alignment with Bowtie 2. Nat Methods 9, 357–359. 10.1038/nmeth.1923.

75. Meers, M.P., Tenenbaum, D., and Henikoff, S. (2019). Peak calling by Sparse Enrichment Analysis for CUT&RUN chromatin profiling. Epigenetics Chromatin 12, 42. 10.1186/s13072-019-0287-4.

76. Ramírez, F., Ryan, D.P., Grüning, B., Bhardwaj, V., Kilpert, F., Richter, A.S., Heyne, S., Dündar, F., and Manke, T. (2016). deepTools2: a next generation web server for deep-sequencing data analysis. Nucleic Acids Res 44, W160–165. 10.1093/nar/gkw257.

77. Bagwell, C.B., Inokuma, M., Hunsberger, B., Herbert, D., Bray, C., Hill, B., Stelzer, G., Li, S., Kollipara, A., and Ornatsky, O. (2020). Automated data cleanup for mass cytometry. Cytometry Part A 97, 184–198.

