## Supplementary figure for "Mutant EZH2 alters the epigenetic network and increases epigenetic heterogeneity in B cell lymphoma"

### Supplementary Fig. 1

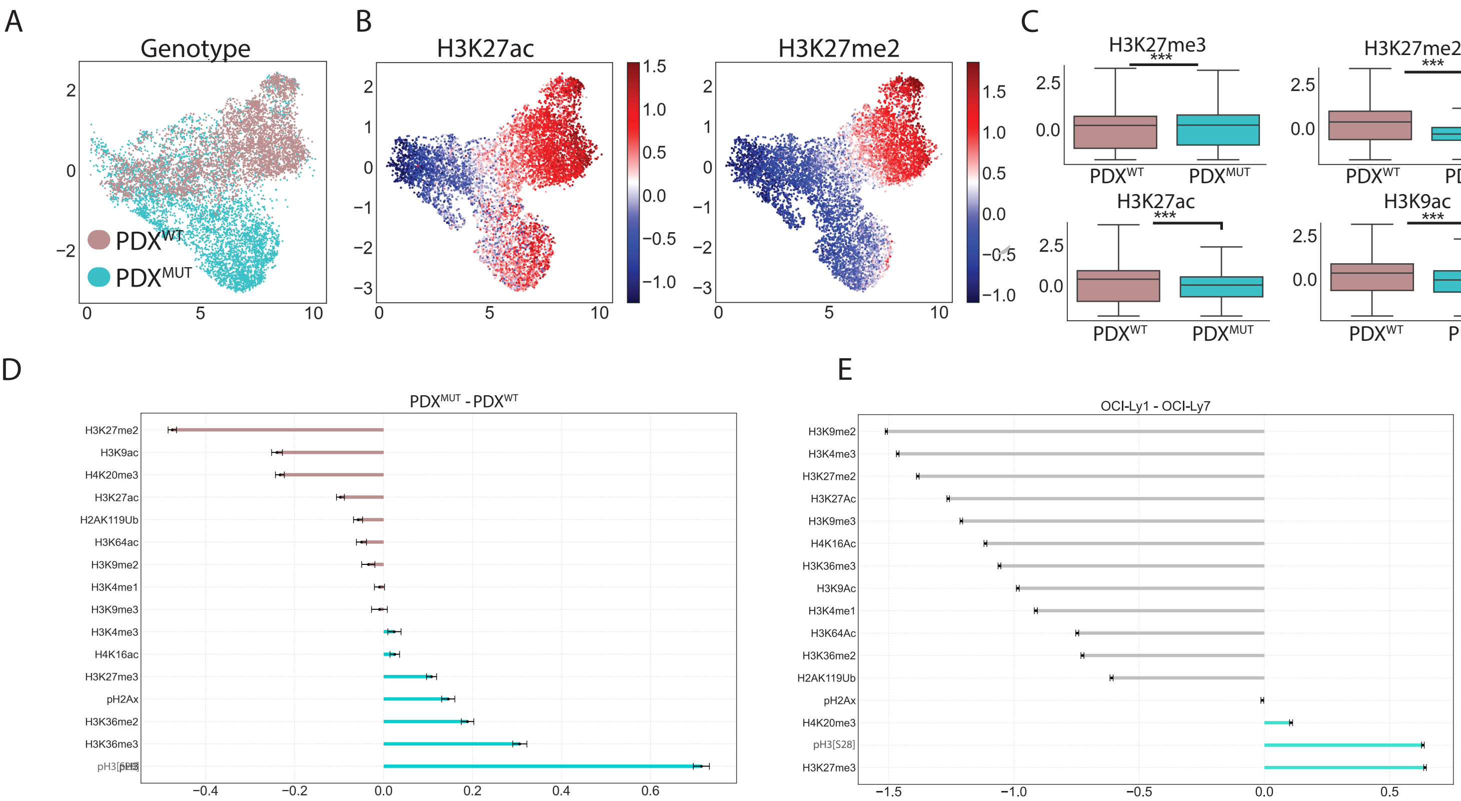

Supplementary Fig. 2

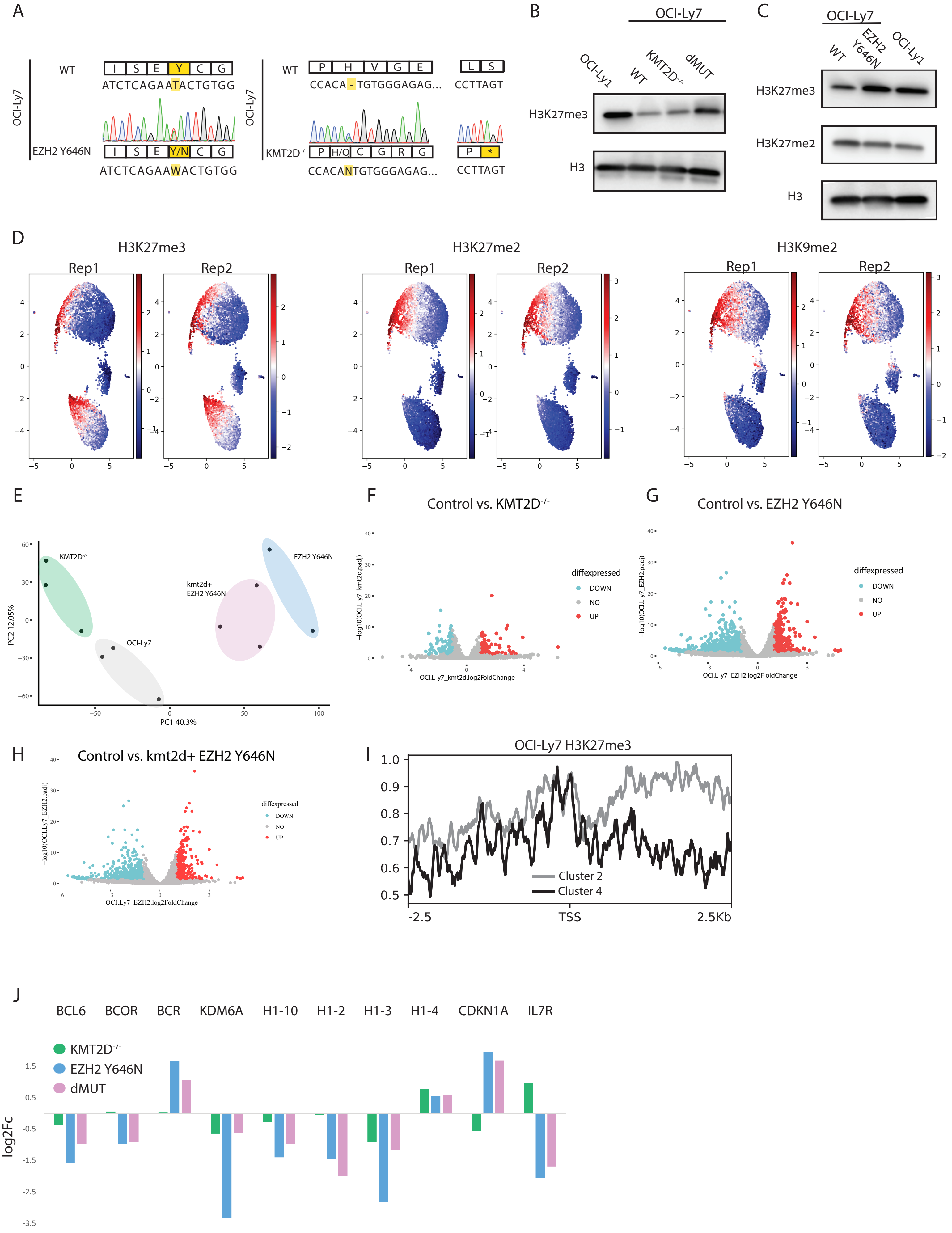

Supplementary Fig. 3

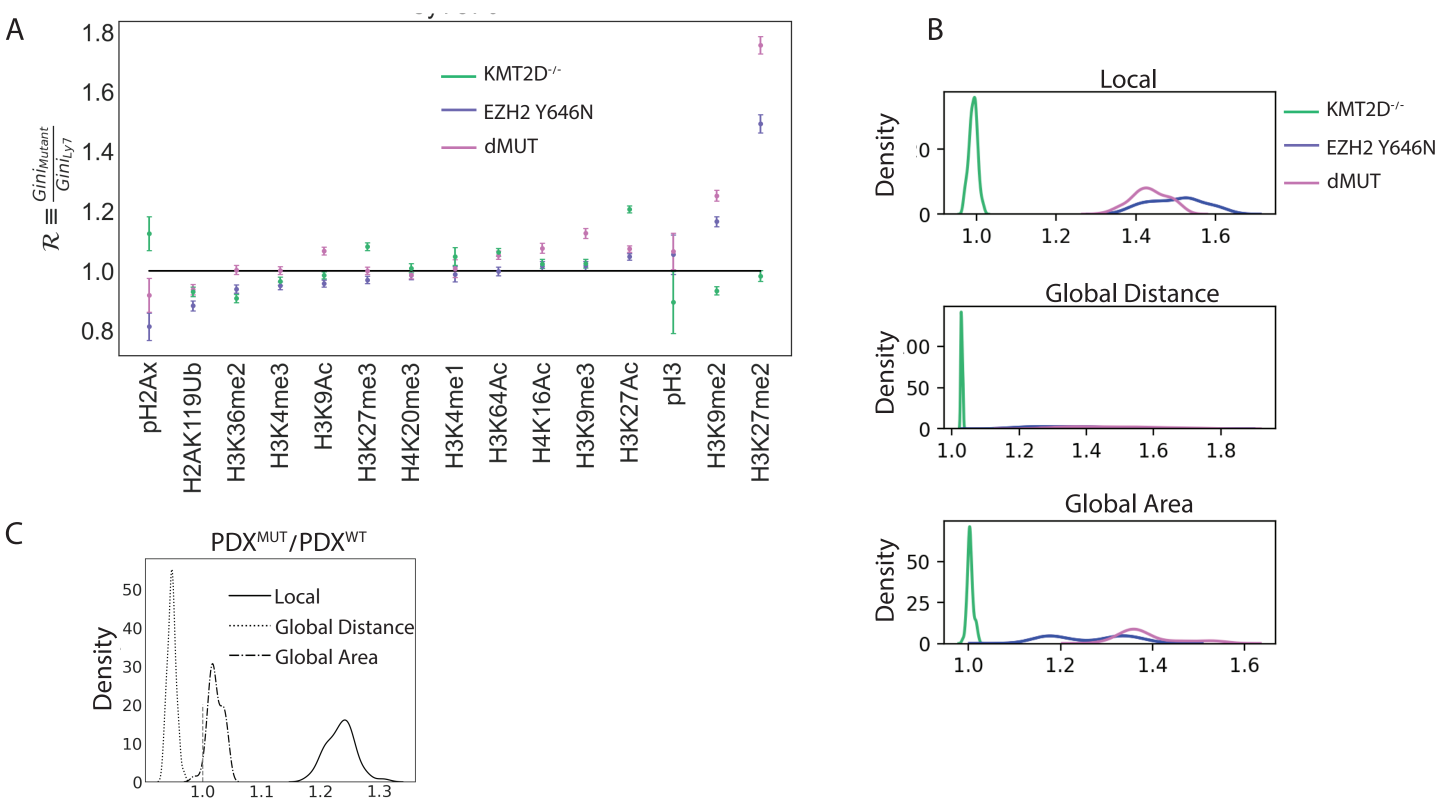

### Supplementary Fig. 4

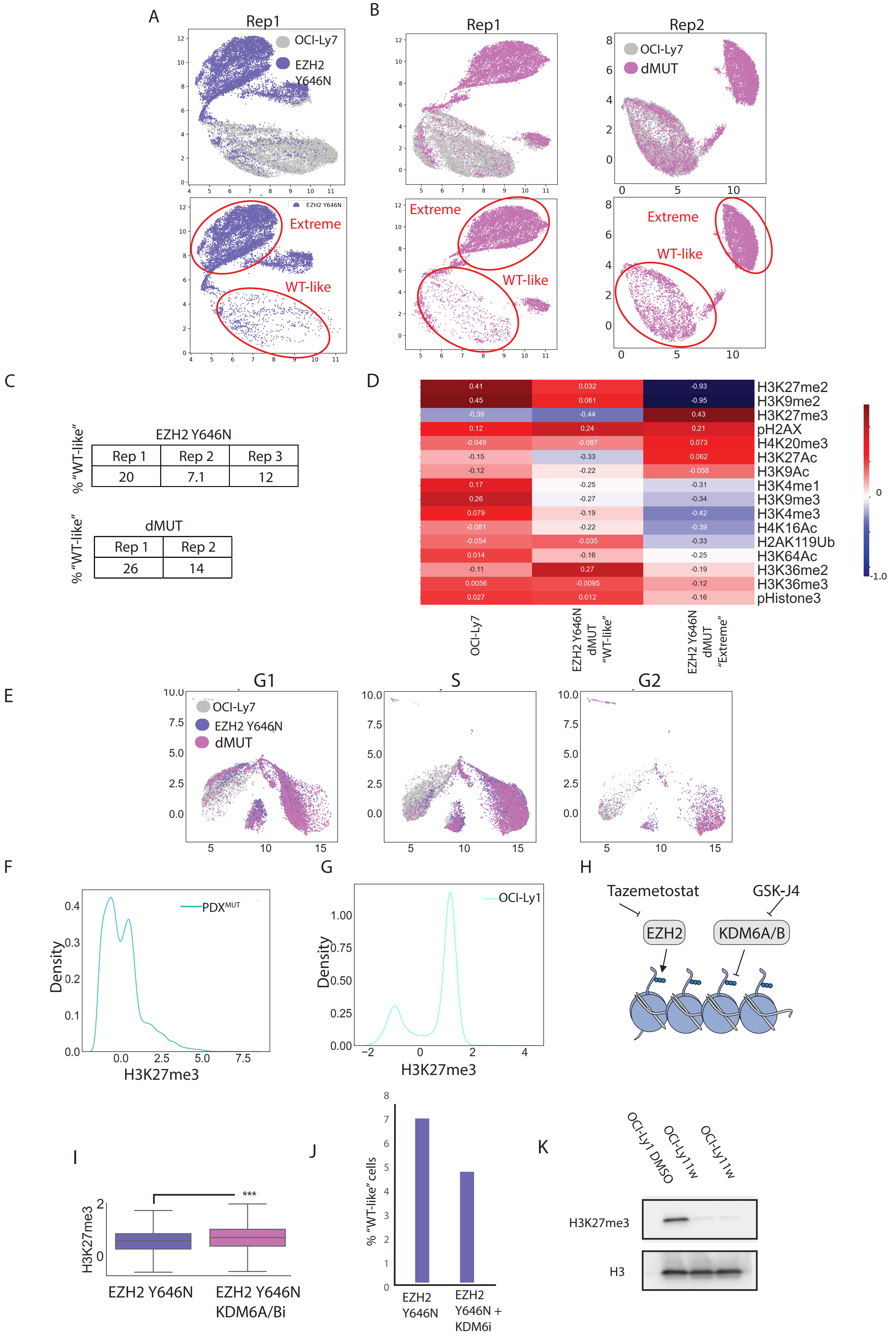

Supplementary Fig. 5

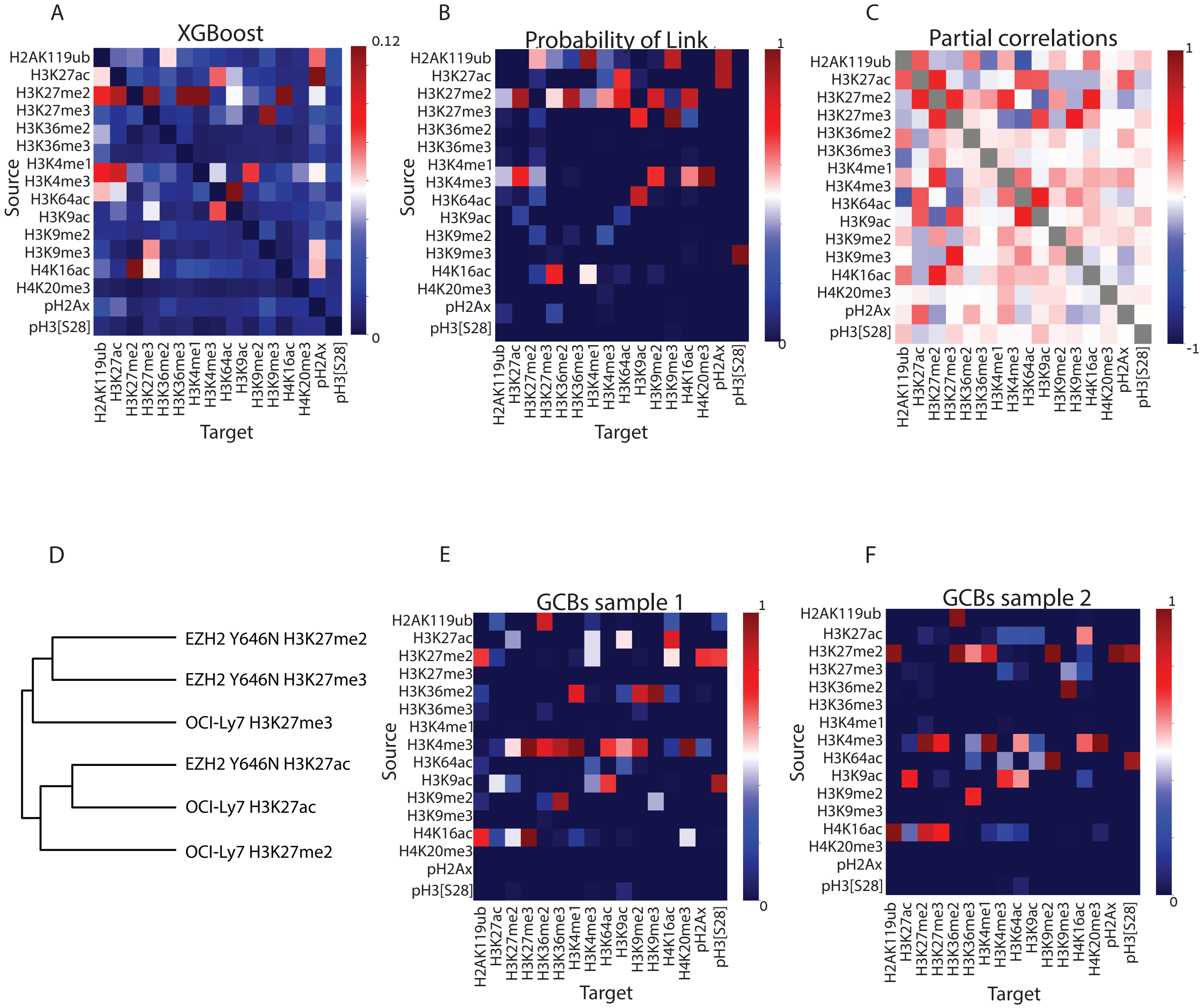

Supplementary Fig. 6

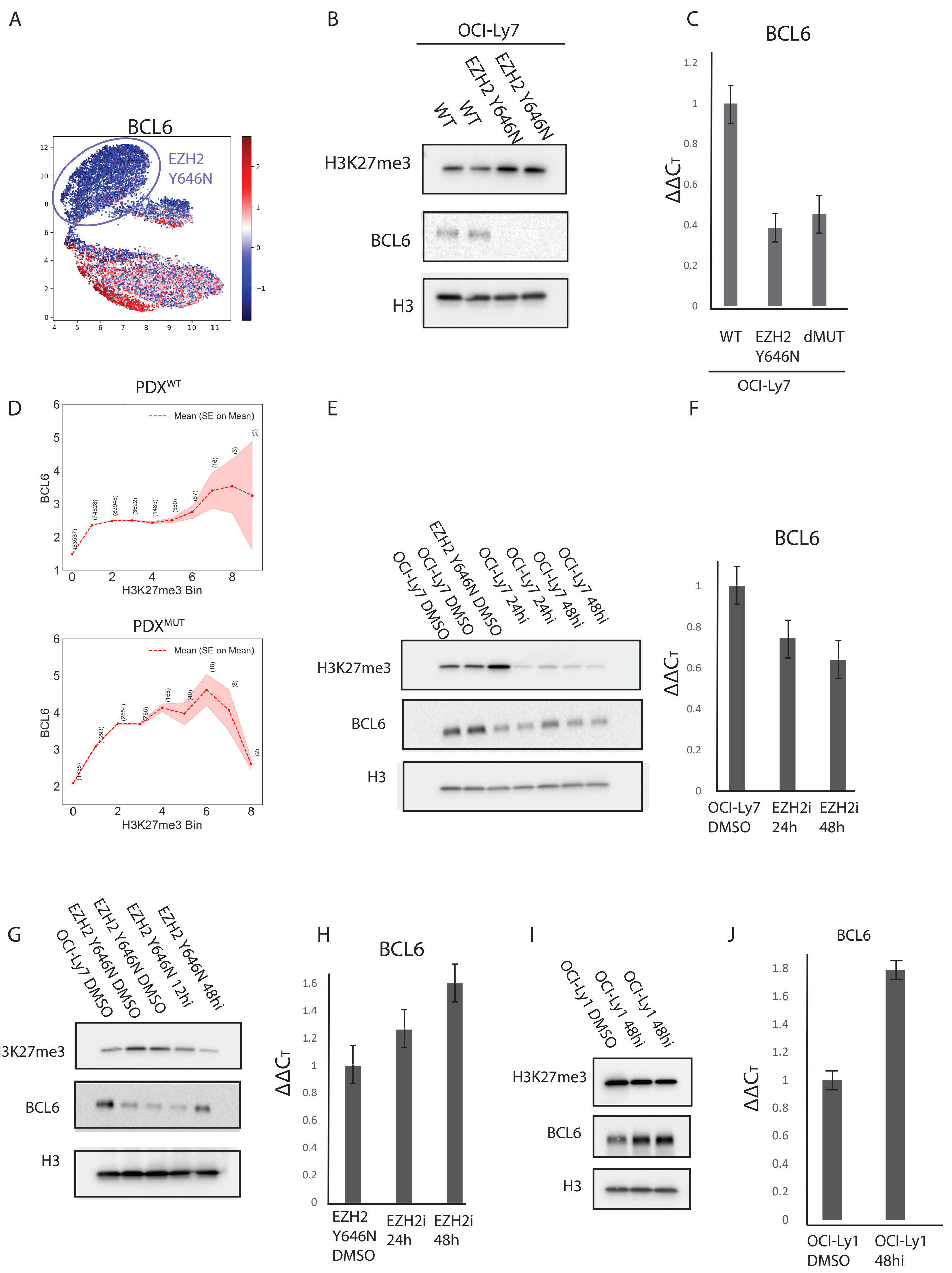
