## Supplementary material for "Mutant EZH2 alters the epigenetic network and increases epigenetic heterogeneity in B cell lymphoma": Figure legends for Supplementary figures

**Supplementary Figure S1**

**A.** Two patient-derived xenografts; PDX^WT^ which carries WT copies of EZH2 and KMT2D, and PDX^MUT^ which carries the EZH2 Y646N mutation and biallelic loss of KMT2D, were analyzed by CyTOF. UMAP was performed based on all epigenetic marks measured, following scaling and normalization. Colors indicate the sample index.

**B.** Scaled, normalized levels of the indicated histone modifications on the UMAP of the PDXs that is shown in A.

**C.** Expression levels of the indicated modifications in PDX^WT^ and PDX^MUT^, as measured by the CyTOF. P values were calculated by Welch’s t test. ***p value < 0.001.

**D.** Mean differences between PDX^WT^ and PDX^MUT^ for the indicated histone modifications. The mean values for PDX^WT^ were subtracted from PDX^MUT^. Uncertainties were estimated using bootstrapping.

**E.** Mean differences between OCI-Ly7 and OCI-Ly1 for the indicated histone modifications. The mean values for OCI-Ly7 (WT) were subtracted from OCI-Ly1 (EZH2 Y646N). Uncertainties were estimated using bootstrapping.

**Supplementary Figure S2**

**A.** Left: Sequencing traces of Exon 16 of EZH2, indicating the heterozygous gain-of-function EZH2 Y646N mutation, generated by CRISPR-Cas9 genome editing, in OCI-Ly7 cells. Right: Sequencing traces of Exon 2 of KMT2D, indicating a biallelic insertion of A/C by CRISPR-Cas9 genome editing resulting in a premature stop codon, in OCI-Ly7 cells.

**B-C.** Western blot analysis of the indicated modifications in the isogenic OCI-Ly7 WT cells and their counterparts carrying the EZH2 GOF mutation, KMT2D knockout, or a combination of both. Also shown are OCI-Ly1 cells expressing mutant-EZH2 and biallelic loss of KMT2D. Histone H3 represents loading control.

**D.** OCI-Ly7 cells (WT), as well as their isogenic counterparts carrying mutant-EZH2 (EZH2 Y646N), biallelic knockout of KMT2D (KMT2D^-/-^), or a combination of both mutations in EZH2 and KMT2D (dMUT), were analyzed by CyTOF. UMAP was performed based on all epigenetic marks measured in two independent biological repeats, following scaling and normalization (see Figure 1E). Shown are the indicated modifications.

**E.** RNA-sequencing analysis of OCI-Ly7 cells and the indicated isogenic mutant lines. Principal component analysis of count number for all genes. Showing PC1 (40.3%) and PC2 (12.05%).

**F-H.** Volcano plots showing Log fold change and p.adj values of differentially expressed genes between the indicated mutant lines versus OCI-Ly7. The list of differentially expressed genes was obtained using DESeq2. Upregulated genes marked in blue were filtered by >1 LogFC and p.adj <= 0.05. Downregulated genes marked in red were filtered by <-1 LogFC and p.adj <= 0.05

**I.**  Visualization of OCI-Ly7 H3K27me3 read coverage around the TSS, as measured by Cut&Run, of differential genes from clusters 2 and 4 (Figure 1H).

**J.** Bar plot showing Log2 change values of the indicated genes in the mutant cell lines versus OCI-Ly7. Colors indicate sample index.

**Supplementary Figure S3**

**A.** The Gini coefficient, commonly used to measure the inequality among the values of a frequency distribution^44^, was used to determine the relative heterogeneity of each modification between the different cell lines. The ratio of the Gini coefficient between the individual lines and the WT was used as a measure of the excess heterogeneity of the mutant over the WT for each modification.

**B.** The indicated heterogeneity measurements that were defined in Figure 2C were calculated for OCI-Ly7 cells versus each of the mutant lines, across a wide range of UMAP parameters. This experiment is a biological repeat of the experiment presented in Figure 2C. Plotted are the ratios of the values of each mutant line versus OCI-Ly7. Thus, values over 1 indicate excess heterogeneity in the mutant over that of the WT. EZH2 mutant cells showed higher heterogeneity in all measurements.

**C.** Heterogeneity measurements calculated for the two patient derived xenografts with WT EZH2 (PDX^WT^) versus mutant-EZH2 (PDX^MUT^). Overall, EZH2-mutant cells show higher heterogeneity locally, and for global area.

**Supplementary Figure S4**

**A-B.** Top: OCI-Ly7 cells expressing WT EZH2, mutant-EZH2 (EZH2 Y646N), or a combination of mutant-EZH2 with KMT2D biallelic knockout (dMUT) were analyzed by CyTOF. Shown are joint UMAPs of OCI-Ly7 cells (WT) and the indicated lines. A represents a repeat of the experiment shown in Figure 2F. B shows two repeats for the double mutant (KMT2D^-/-^ EZH2 Y646N). Colors indicate the sample index. Bottom: Only mutant-EZH2 cells are plotted, to highlight the subpopulation of mutant cells that cluster with WT cells, referred to as ‘WT-like’.

**C.** The fraction of WT-like cells in three biological CyTOF replicates for the single EZH2 Y6464 mutant line, and two biological replicates for the double mutant. The percentage of cells consisting of this subpopulation is dynamic and varies between experiments.

**D.** The mean of distribution of the indicated histone modifications for OCI-Ly7 cells expressing WT EZH2 (‘WT’), and for the double mutant of EZH2 Y646N with KMT2D knockout (dMUT) cells that either express robustly the GOF phenotype and form a distinct cluster (EZH2 Y646N dMUT ‘Extreme’), or the ‘WT-like’ subpopulation that clusters with OCI-Ly7 WT cells. See also Figure 2G for the single EZH2 mutant line.

**E.** OCI-Ly7 cells and the indicated isogenic mutants were analyzed by CyTOF, that included all epigenetic modifications as well as the Maxpar cell cycle panel kit. The cell cycle markers, included in the CyTOF panel, were used to determine the cell cycle phase of each cell: G1, S and G2. For each phase, a joint UMAP of the WT and mutant cells was generated, based on all epigenetic modifications. ‘WT-like’ cells are observed for each of the indicated cell cycle phases.

**F-G.** Histogram of H3K27me3 levels in: **F.** Patient-derived xenograft expressing mutant-EZH2 (PDX^MUT^). **G.** OCI-Ly1 cells expressing mutant-EZH2. Cells expressing the mutant enzyme show bimodal distribution of H3K27me3, indicating two distinct subpopulations.

**H.** Illustration of the effects of EZH2 inhibitor (EZH2i) Tazemetostat that blocks EZH2-mediated deposition of H3K27me2/3, and GSK-J4 (KDM6i) that inhibits the H3K27 demethylase KDM6A/B, resulting in elevated H3K27me3 levels.

**I.** OCI-LY7 with mutant-EZH2 were treated with the histone lysine 27 demethylase KDM6A/B inhibitor GSK-J4 at a concentration of 1µM for 24 hours, or left untreated as control. Shown are the expression levels of H3K27me3, as measured by CyTOF. Treatment with the inhibitor led to the expected upregulation of H3K27me3 levels. P values were calculated by Welch’s t test. ***p value < 0.001.

**J.** OCI-Ly7 cells were treated with KDM6A/B inhibitor at 1µM concentration for 24 hours, followed by CyTOF. The percentage of ‘WT-like’ cells in control and EZH2i-treated cells is shown. EZH2 inhibition reduced the fraction of ‘WT-like’ cells.

**K.** OCI-Ly1 cells expressing mutant-EZH2 were treated with EZH2 inhibitor at a concentration of 10µM for one week to completely deplete H3K27me3. Shown is a western blot for H3K27me3 in the control and two repeats of the treated cells. Histone H3 is used as a loading control.

**Supplementary Figure S5**

**A-C**. Models to decipher interactions within the epigenetic network, applied to unperturbed OCI-Ly7 single-cell CyTOF data, on a biological repeat of the experiment shown in Figure 3A-C. **A.** XGBoost analysis. **B.** ‘Probability of link’ analysis. For both A and B, the Y axis indicates ‘source’ modification and X axis indicates its ‘target’ modification affected by the Y axis. **C.** Partial correlations between histone modifications.

**D.** Dendrogram of the hierarchical clustering of spearman correlations of Cut&Run normalized reads between all samples, calculated on 10kbp genomic bins.

**E-F.** Probability of link analysis done on CyTOF data of unperturbed germinal-center B cells (CD20^+^, CD38^+^), derived from tonsils, for two independent samples derived from different patients. Tonsils were dissociated to single cells followed by staining with the panel of metal-conjugated antibodies and CyTOF analysis.

**Supplementary Figure S6**

**A.** Scaled, normalized levels of BCL6 on the joint UMAP of OCI-Ly7 and EZH2 Y646N cells, corresponding to the UMAP shown in Figure 2A. Cells expressing mutant-EZH2 show downregulation of BCL6 levels.

**B.** Western blot analysis of H3K27me3 and BCL6 in the indicated samples. Histone H3 is used as a loading control.

**C.** Quantitative RT-PCR analysis of BCL6 expression in the isogenic OCI-Ly7 WT and EZH2-mutant cells. ΔΔC_T_ values relative to OCI-Ly7 ±s.d (n=3) are shown. HPRT was used for normalization.

**D.** BCL6 mean levels in cells binned according to H3K27me3 levels. Red hue represents standard error of the mean. Number of cells in each bin is shown. Top: PDX with WT-EZH2 (PDX^WT^). Bottom: PDX with mutant-EZH2 (PDX^MUT^). H3K27me3 and BCL6 show non-linear relationship.

**E-F.** Western blot analysis of H3K27me3 and BCL6. Histone H3 is used as a loading control. **E.** OCI-Ly7 cells were treated with EZH2 inhibitor at a concentration of 10µM for the indicated times. **F.** EZH2-mutant cells were treated with EZH2 inhibitor at a concentration of 10µM for 48 hours.

**G-H.** Quantitative RT-PCR analysis of BCL6 expression in the isogenic OCI-Ly7 WT and EZH2-mutant cells, treated with EZH2i 10µM at the indicated times. ΔΔC_T_ values relative to OCI-Ly7 ±s.d (n=3) are shown. HPRT was used for normalization.

**I.** Western blot analysis of H3K27me3 and BCL6. OCI-Ly1 cells, carrying mutant-EZH2, were treated with EZH2 inhibitor at a concentration of 10µM for 48 hours.

**J.** Quantitative RT-PCR analysis of BCL6 expression in OCI-Ly1 cells treated with EZH2 inhibitor at a concentration of 10µM for 48 hours versus DMSO control. ΔΔC_T_ values relative to DMSO treated sample ±s.d (n=3) are shown.

**References**

44.	Gini, C. (1936) On the measure of concentration with espacial reference to income and wealth. *Cowles Commission*, **2**.
